# Diurnal variation in xylem water isotopic signature biases depth of root-water uptake estimates

**DOI:** 10.1101/712554

**Authors:** Hannes De Deurwaerder, Marco D. Visser, Matteo Detto, Pascal Boeckx, Félicien Meunier, Liangju Zhao, Lixin Wang, Hans Verbeeck

**Affiliations:** CAVElab - Computational & Applied Vegetation Ecology, Faculty of Bioscience Engineering, Ghent University, Ghent, Belgium; Department of Ecology and Evolutionary Biology, Princeton University, Princeton, NJ, USA; ISOFYS – Isotope Bioscience Laboratory, Faculty of Bioscience Engineering, Ghent University, Ghent, Belgium; Ecological Forecasting Lab, Department of Earth and Environment, Boston University, Boston, Massachusetts, USA; Shaanxi Key Laboratory of Earth Surface System and Environmental Carrying Capacity, College of Urban and Environmental Sciences, Northwest University, Xi’an 710127, China; Key Laboratory of Ecohydrology and Integrated River Basin Science, Northwest Institute of Eco-Environment and Resources, Chinese Academy of Sciences, Lanzhou 730000, China; Department of Earth Sciences, Indiana University-Purdue University Indianapolis (IUPUI), Indianapolis, IN 46202, USA

**Keywords:** Deuterium, Ecohydrology, Lianas, Root water uptake, Sap flow, Stable isotopes, Tropical trees, Water competition

## Abstract

- Stable water isotopes are a powerful and widely used tool to derive the depth of root water uptake (RWU) in lignified plants. Uniform xylem water isotopic signature (*i-H*_*2*_*O-xyl*) along the length of a lignified plant is a central assumption, which has never been properly evaluated.
- Here we studied the effects of diurnal variation in RWU, sap flow velocity and various other soil and plant parameters on *i-H*_*2*_*O-xyl* signature within a plant using a mechanistic plant hydraulic model.
- Our model predicts significant variation in *i-H*_*2*_*O-xyl* along the full length of an individual plant arising from diurnal RWU fluctuations and vertical soil water heterogeneity. Moreover, significant differences in *i-H*_*2*_*O-xyl* emerge between individuals with different sap flow velocities. We corroborated our model predictions with field observations from French Guiana and northwestern China. Modelled *i-H*_*2*_*O-xyl* varied considerably along stem length ranging up to 18.3‰ in δ^2^H and 2.2‰ in δ^18^O, largely exceeding the range of measurement error.
- Our results show clear violation of the fundamental assumption of uniform *i-H*_*2*_*O-xyl* and occurrence of significant biases when using stable isotopes to assess RWU. As a solution, we propose to include monitoring of sap flow and soil water potential for more robust RWU depth estimates.

## Introduction

The use of stable water isotopes has greatly enhanced ecohydrology studies by providing insights into phenomena that are otherwise challenging to observe, such as depth of root water uptake (RWU) (Rothfuss & Javaux, 2017), below ground water competition and hydraulic lift (Hervé-Fernández *et al.*, 2016; Meunier *et al.*, 2017). Compared to root excavation, the technique is non-destructive, far less labor-intensive and informs on actual RWU while excavation solely informs on root distribution and architecture. Moreover, its flexibility allows use across multiple scales both spatial (i.e. individual to ecosystem) and temporal (i.e. daily to seasonal; Dawson *et al.* 2002). The advantages and wide applicability of this method have it a popular technique that pushes the boundaries of ecohydrology (Dawson *et al.*, 2002; Yang *et al.*, 2010; Rothfuss & Javaux, 2017).

A variety of methods exist that infer RWU depth from plant isotopic signatures, but all rely on a direct relationship between the isotopic compositions of plant xylem water and soil water (Ehleringer & Dawson, 1992). More precisely, all have two key assumptions. The first is that water isotopic signature remains unchanged during transport from root uptake to evaporative sites (e.g. leaves and non-lignified green branches). Hence, isotope fractionation - processes that shift the relative abundance of the water isotopes during root water uptake and water through non-evaporative tissues - is neglected (Wershaw *et al.*, 1966; Zimmermann *et al.*, 1967; White *et al.*, 1985; Dawson & Ehleringer, 1991; Walker & Richardson, 1991; Dawson *et al.*, 2002; Zhao *et al.*, 2016). Second, all methods assume that xylem water provides a well-mixed isotopic signature of water from different soil layers: sampled xylem water instantaneously reflects the distribution and water uptake of the roots independent of sampling time or height.

The first assumption is relatively well supported. Fractionation at root level has not raised concerns for most RWU assessments using water isotopes (Rothfuss & Javaux, 2017), with the exception of kinetic fractionation. Kinetic fractionation is a process driven by the differences in molecule mass among the isotopologues that occurs only in extreme environments (Lin & Sternberg, 1993; Ellsworth and Williams, 2007; Zhao *et al.*, 2016). Similarly, isotopic fractionation of water within an individual plant, although possible, is not considered a serious problem (Yakir, 1992; Dawson & Ehleringer, 1993; Cernusak *et al.*, 2005; Mamonov *et al.*, 2007; Zhao *et al.*, 2016). However, the second assumption of time and space invariance of xylem isotopic water signatures has, to our best knowledge, never been assessed

In principle, temporal variance in xylem water isotopic signature within a plant during a day or along its height can be expected on first principles. Here we hypothesize that it is in fact likely that various plant physiological processes, ranging from very simple to more complex mechanisms could influence within plant variance in *i-H*_*2*_*O-xyl* at short time scales. For instance, plant transpiration during the course of the day is regulated by atmospheric water demand and leaf stomata which have clear and well known diurnal patterns (Steppe & Lemeur, 2004; Epila *et al.*, 2017). This results in changing water potential gradients within the soil-plant-atmosphere continuum and therefore fluctuations in the depth RWU are also expected (Goldstein *et al.*, 1998; Doussan *et al.*, 2006; Huang *et al.*, 2017). Hence, as we expect plants capacity to take up water at different soil layers to shift during the day, we should also expect diurnal variation in the mixture of water isotopes taken up from various depths. As water moves up along the xylem with velocity proportional to sap flow, different plants and species might respond differently to diurnal variation in RWU. Therefore, from very basic principles we may expect temporal variation in xylem water isotopic signature (*i-H*_*2*_*O-xy)* to propagate to different plant heights. As sap flow velocity depends on plant hydraulic traits in relation to atmospheric water demand and soil moisture gradient, this mechanism could make comparison of isotopic data among individuals and species misleading.

In this study we provide a critical assessment of the assumption of *i-H*_*2*_*O-xyl* invariance over time and along the length of plant stems. We test the hypothesis that major alterations in the *i-H*_*2*_*O-xyl* along the length of lignified plants arise naturally during the day and that this variation in *i-H*_*2*_*O-xyl* exceeds the expected measurement error. We test this hypothesis with a twofold approach. First, we build a simple mechanistic model that incorporates basic plant hydraulic realism. We use this model to specifically test that even rudimentary mechanistic models of plant hydraulic functioning predict that diurnal changes in the soil-plant-atmosphere continuum result in shifting mixtures of soil isotopes absorption Second, we test whether the *i-H*_*2*_*O-xyl* sampled at different plant heights or at different times of the day show large variances with field observations from i) six Neotropical canopy trees and six Neotropical canopy lianas sampled at different heights in French Guiana, and ii) high temporal resolution *i-H*_*2*_*O-xyl* data of 6 distinct plant species from the Heihe River Basin in northwestern China (Zhao *et al.*, 2014).

## Materials and Methods

### Part A: Modelling exploration

#### Model derivation

The expected *i-H*_*2*_*O-xyl* at different stem heights within a tree during the course of the day can be derived from plant and physical properties such as root length density, total fine root surface area, water potential gradients and soil water isotopic signatures. We call this the SWIFT model (i.e. <u>S</u>table <u>W</u>ater Isotopic <u>F</u>luctuation within <u>T</u>rees). To derive the SWIFT model, we first describe the establishment of *i-H*_*2*_*O-xyl* entering the tree at stem base via a multi-source mixing model (Phillips & Gregg, 2003). We subsequently consider vertical water transport within the tree, driven by the established sap flow pattern. Note that the model presented here, focusses on deuterium but can easily be used to study stable oxygen isotopologues. To ensure consistency and clarity in variable declarations we maintain the following notation in the subscripts of variables: uppercase roman to distinguish the medium through which water travels (X for xylem, R for root, S for soil) and lowercase for units of time and distance (*h* for stem height, *t* for time and *i* for soil layer index). A comprehensive list of variables, definitions and units is given in Table 1.

**Table 1.** Nomenclature.

| Symbol | Description | Unit |
| --- | --- | --- |
| $A_{R,i}$ | is the fine root surface area distributed over soil layer $i$. | $\text{m}^2$ |
| $A_{Rtot}$ | The plants' total active fine root surface area | $\text{m}^2$ |
| $A_{SAPWOOD}$ | Sapwood area | $\text{m}^2$ |
| $A_x$ | Total lumen area | $\text{m}^2$ |
| $b$ | Shape parameter for the soil hydraulic properties (Clapp & Hornberger, 1978) | dimensionless |
| $B_i$ | The overall root length density distribution per unit of soil, not necessarily limited to the focal plant. | $\text{m m}^{-3}$ |
| $\delta^2H_{X,0,t}$ | Xylem water isotopic signature of an individual plant at stem base at time $t$ | in ‰ VSMOW |
| $\delta^2H_{X,h,t}$ | Xylem water isotopic signature of an individual plant at height $h$ and time $t$ | in ‰ VSMOW |
| $\delta^2H_{S,i}$ | Soil water isotopic signature of the $i^{\text{th}}$ soil layer (constant over time) | in ‰ VSMOW |
| $\delta_{\text{sample}}$ | Isotopic signature of a sample | in ‰ VSMOW |
| $\Delta\hat{\Psi}_{i,t}$ | Estimated water potential gradient between stem base and the $i^{\text{th}}$ soil layer at time $t$ derived from Eq. (8) | m |
| $\Delta\Psi_{i,t}$ | Soil water potential gradient between soil and roots at the $i^{\text{th}}$ soil layer at time $t$ | $\text{m H}_2\text{O}$ |
| $\epsilon^2\text{H}_X ; \epsilon^{18}\text{O}_X$ | Normalized isotopic water signatures | in ‰ VSMOW |
| $f_{i,t}$ | Fraction of water taken up in the $i^{\text{th}}$ soil layer at time $t$ | dimensionless |
| $h$ | Measurement height | m |
| $i$ | Soil layer index | dimensionless |
| $i\text{-H}_2\text{O-xyl}$ | Xylem water isotopic signature | in ‰ VSMOW |
| $k_i$ | Soil-root conductance of the $i^{\text{th}}$ soil layer | $\text{s}^{-1}$ |
| $K_{max}$ | Maximum soil hydraulic conductivity | $\text{m s}^{-1}$ |
| $k_R$ | Effective root radial conductivity | $\text{s}^{-1}$ |
| $k_S$ | The conductance associated with the radial water flow between the soil and the root surface | $\text{s}^{-1}$ |
| $K_{S,i}$ | Soil hydraulic conductivity at the $i^{\text{th}}$ soil layer | $\text{m s}^{-1}$ |
| $\ell$ | The approximated radial pathway length of water flow between bulk soil and root surface | $\text{m}$ |
| LF | Lumen fraction per unit sapwood area | $\text{m}^2 \text{m}^{-2}$ |
| n | Number of unique contributing water sources | # |
| $\Psi_{sat}$ | Soil water potential at soil saturation | $\text{m}$ |
| $\Psi_{S,i,t}$ | Soil water potential of the $i^{\text{th}}$ soil layer at time $t$ | $\text{m}$ |
| $\Psi_{X,0,t}$ | Water potential at the base of the plant stem at time $t$ | $\text{m}$ |
| R | Heavy to light isotope ratio measured in the sample or standard | % |
| $RWU_{i,t}$ | Net amount of water entering and leaving the root tissues per unit of time in the $i^{\text{th}}$ soil layer at time $t$ | $\text{m}^3 \text{s}^{-1}$ |
| $SF_t$ | Instantaneous sap flow at time $t$ | $\text{m}^3 \text{s}^{-1}$ |
| $SF_S$ | Sap flow velocity, calculated as the sap flow per sapwood area | $\text{m h}^{-1}$ |
| $SF_V$ | True sap flow velocities, calculated as the sap flow per lumen area | $\text{m h}^{-1}$ |
| $\tau$ | Delay before the isotopic signature at stem base reaches stem height $h$ | $\text{s}$ |
| $\theta_{sat}$ | Soil moisture content at soil saturation | $\text{m}^3 \text{m}^{-3}$ |
| $\theta_{S,i,t}$ | Soil moisture content of the $i^{\text{th}}$ soil layer at time $t$ | $\text{m}^3 \text{m}^{-3}$ |
| $z_i$ | Soil depth of the $i^{\text{th}}$ soil layer | $\text{m}$ |

##### Xylem water isotopic signature at stem base

The xylem water deuterium signature (*δ*^2^*H*_*X*,0,*t*_) *of an individual plant at stem base* (i.e. height zero; *h* = 0 m) at time *t*, can theoretically be derived using the multi-source mixing model approach introduced by Phillips & Gregg (2003). Considering a root zone divided into *n* discrete soil layers of equivalent thickness Δ*z*, if the deuterium isotopic signature (*δ*^2^*H*_*S,i*_) of each soil layer is constant over time, a reasonable assumption if the isotopic measurements are conducted during rain-free periods, *δ*^2^*H*_*X*,0,*t*_ can be expressed as: 

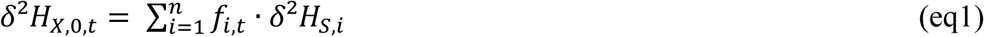

where *f*_*i,t*_ is the fraction of water taken up at the *i*^th^ soil layer defined as: 

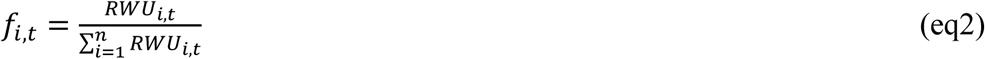

and *RWU*_*i,t*_ is the net amount of water entering and leaving the roots at time *t* in the *i*^th^ soil layer (*RWU*_*i,t*_ is defined positive when entering the root). The current representation of the model assumes no water loss via the root system and no mixing of the extracted water from different soil layers within the roots until the water enters the stem base. When tree capacitance is neglected, the sum of *RWU*_*i,t*_ across the entire root zone is equal to the instantaneous sap flow at time *t, SF*_*t*_.: 

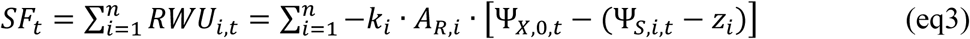

Where *k*_*i*_ is the plant specific total soil-to-root conductance, *Ψ*_*X*,0,*t*_ is the water potential at the base of the plant stem and *Ψ*_*S,i,t*_ is the soil water matric potential. Total plant water potential is generally defined as the sum of the pressure, gravity and matrix potential. Hence, Ψ_*X*,0,*t*_ represents the xylem pressure potential. The term *z*_*i*_ is the gravimetric water potential necessary to lift the water from depth *z*_*i*_ to the base of the stem, assuming a hydrostatic gradient in the transporting roots. The model considers *z*_*i*_ to be a positive value (zero at the surface), thus *z*_*i*_ is subtracted from Ψ_*S,i,t*_. A_R,i_ is the fine root surface area distributed over soil layer *i*. This parameter can be derived from plant allometric relations (Čermák *et al.*, 2006) which is subsequently distributed over the different soil layers via Jackson *et al.* (1995).

The total soil-to-root conductance is calculated assuming the root and soil resistances are connected in series: 

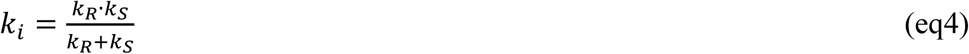

where *k*_*R*_ is the effective root radial conductivity (assumed constant and uniform), and *k*_*S*_ = *K*_*S,i*_/*𝓁* is the conductance associated with the radial water flow between soil and root surface. 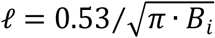 *B*_*i*_ represents the effective radial pathway length of water flow between bulk soil and root surface (Vogel *et al.*, 2013). *B*_*i*_ represents the overall root length density distribution per unit of soil. *K*_*S,i*_ is the soil hydraulic conductivity for each soil depth. *K*_*S,i*_ depends on soil water moisture and thus relates to the soil water potential *Ψ*_*S,i,t*_ of the soil layer where the water is extracted. *K*_*S,i*_ is computed using the Clapp & Hornberger (1978) formulation: 

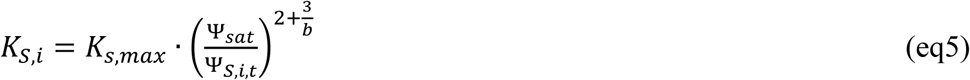

where *K*_*s,max*_ is the soil conductivity at saturation and *b* and Ψ_*sat*_ are empirical constants that depend on soil type (here considered as constant through all soil layers).

Subsequently, *f*_*i*.*t*_ can be restructured as: 

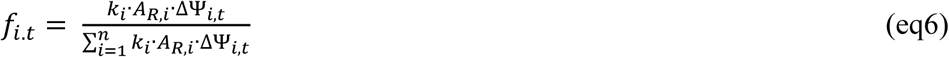

where the root to soil water potential gradient is represented as ΔΨ_*i,t*_ = Ψ_*X*,0,*t*_ − (Ψ_*S,i,t*_ − *z*_*i*_). Combining eq.(1) and eq.(6) then allows derivation of *δ*^2^*H*_*X*,0,*t*_ as follows: 

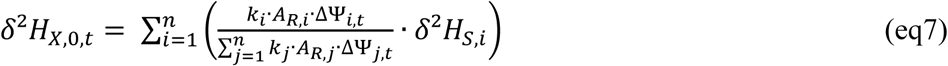

This equation requires estimates of ΔΨ_*i,t*_, which is preferably measured instantaneously in the field (i.e. via stem and soil psychrometers for Ψ_*X*,0,*t*_ and Ψ_*S,i,t*_, respectively). However, as measurements of Ψ_*X*,0,*t*_ are not always available, estimated 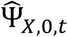 can be derived from sap flow by re-organizing eq.(3) into: 

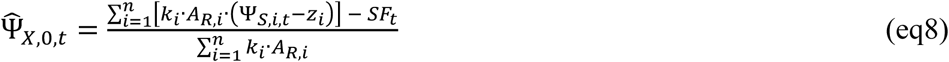

which then allows replacement of Ψ_*X*,0,*t*_ with 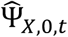 in eq.(7).

##### Height-dependent xylem water isotopic signature

In our model, from the stem base, the water isotopologues simply move upwards with the xylem sap flow, hence diffusion and water fractionation during transportation are not considered. The xylem water isotopic signature at height *h* and time *t* (*δ*^2^*H*_*X,h,t*_) is then the water isotopic signature at stem base at time *t* – τ. 

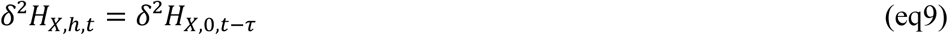

where *τ* is the lag before *δ*^2^*H*_*X*,0,*t*_ reaches stem height *h* which depends only on the true sap flow velocity in the xylem (*SF*_*V*_). True sap flow velocity indicates the real speed of vertical water displacement within a plant, derived by dividing *SF*_*t*_ over the lumen area of the plant (*A*_*x*_), i.e. the total cross-sectional area of the vessels. *τ* was derived from the mass conservation equality: 

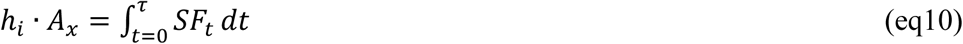

Note that since most scientific studies express sap flow velocity as the sap flow over the total sapwood area (*SF*_*S*_), rather than over the lumen area (*SF*_*V*_), for consistency, we will present the model outputs as functions of *SF*_*S*_.

#### Model parameterization and analyses

We adopted the basic plant parameters from Huang *et al.* (2017) for a loblolly pine (*Pinus taeda L.*) (Table S1). We started with synthetic sap flow patterns and volumes extracted from the model runs of Huang *et al.* (2017) for a typical day (day 11 of the 30 days sequence), and assumed no variation between days. Sap flow follows the plant’s water demand which is the result of daily cycles of transpiration driven by photosynthetic active solar radiation (PAR), vapor pressure deficit (VPD) and optimal stomatal response (Epila *et al.*, 2017). Secondly, both the soil water potential (*Ψ*_*S,i,t*_) and soil water deuterium signature (*δ*^2^*H*_*S,i*_) profiles with soil depth were adopted from Meißner *et al.* (2012) (Fig S2, see Table S1 for equations) and were assumed to stay constant over time. Since measurements of Meißner *et al.* (2012) are derived from a silt loam plot in the temperate climate of central Germany, soil parameters were selected accordingly from Clapp & Hornberger (1978). Subsequently, the following model simulations were executed:

1. **Analysis A1: impact of temporal SF**_**t**_ **variation on isotopic signature at a fixed stem height.** Temporal patterns in xylem deuterium isotopic signatures (*δ*^2^*H*_*X*_) were evaluated for a typical situation, i.e. measurement at breast height (*h*=1.30 m), conforming to standard practice of RWU assessment.
2. **Analysis A2: impact of temporal SF**_**t**_ **variation at different tree heights.** Temporal patterns in *δ*^2^*H*_*X,i*_ within a tree at various sampling heights (5, 10 and 15 m).
3. **Analysis A3: impact of temporal SF**_**t**_ **variation on isotopic signature and the timing of sampling.** Representation of the profile of *δ*^2^*H*_*X*_ along the full height of a tree, measured at different sampling times (9:00, 11:00 and 13:00), with the standard parameterization given in Table S1.
4. **Analysis B: variation in *δ***^2^***H***_***X***_ **due to differences in absolute daily average sap flow speed.** Diurnal patterns in xylem water deuterium signatures in trees that differ solely in daily averaged *SF*_*V*_, which are set to 0.56, 0.28 and 0.14 m h^−1^ (respectively corresponding to *SF*_*S*_ values of 0.08, 0.04 and 0.02 m h^−1
^). All parameters (e.g. RWU) of the four analyses are given in Table S1.

Model runs for each analyses were compared to a null model. The null model adopts the standard assumption of zero variation in *δ*^2^*H*_*X*_ along the length of the plant body. We used extraction protocol related measurement errors with an accepted maximum error range of 3‰ when water extraction recovery rates are higher than 98% (Orlowski *et al.*, 2013). In our null model, this is represented by a normal distribution with a mean of 0‰ and a standard deviation of 1‰, i.e. N(μ=0‰, σ=1‰), which makes the probability of an error of ≥3‰ highly unlikely (p ≤ 0.0027). Analytical errors introduced by the measurement device, i.e. a Picarro (California, USA), are considered negligible relative to the extraction error. Note that *SF*_*V*_, which normalizes sap flow over lumen area, is correlated with plant DBH which enables comparison with field measurements without the need for explicit consideration of DBH in the model. SWIFT was implemented in R version 3.4.0 (R Core Team, 2017), and is publicly available (see GitHub repository HannesDeDeurwaerder/SWIFT).

#### Estimation of rooting depth

RWU depths were derived from the simulated *δ*^2^*H*_*X*_ values by use of both the direct inference method and the end-member mixing analysis method, together representing 96% of the applied methods in literature (Rothfuss & Javaux, 2017). We refer readers to Rothfuss & Javaux (2017) for a complete discussion of these techniques. Here, average rooting depth is assumed to be the depth obtained by relating the simulated *δ*^2^*H*_*X*_ with the *δ*^2^*H*_*S,i*_ depth profile. We compared rooting depth estimates from simulated *δ*^*2*^*H*_*X*_, as described in the analyses above, with the true average rooting depth. The true average rooting depth was defined as the depth corresponding to the daily weighted average *δ*^*2*^*H*_*X*_, calculated as the weighted sum of *δ*^2^*H*_*X,i,t*_ and the relative fraction of water taken up at each depth.

#### Sensitivity analysis

We performed two sensitivity analyses to assess the relative importance of all parameters in generating variance in *δ*^*2*^*H*_*X*_ along the length of a plant. In both sensitivity analyses, we varied model parameters one-at-the-time to assess the local sensitivity of the model outputs. The sensitivity analysis provides insight into the design of field protocols, revealing potential key measurements in addition to any caveats.

We first assessed model sensitivity to (bio)physical variables by modifying model parameters of soil type, sap flow and root properties as compared to the standard parameterization (given in Table S1). The following sensitivity analyses were evaluated:

##### Soil type

The soil moisture content over all soil layers (θ_*S,i,t*_) can be deduced from the *Ψ*_*S,i,t*_ profile using the Clapp & Hornberger (1978) equation: 

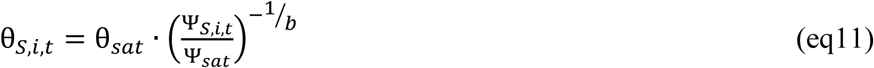

Where θ_*sat*_, Ψ_*sat*_ and *b* are soil-type specific empirical constants that correspond to sandy loam soil textures in the standard model parameterization (Clapp & Hornberger, 1978). The derived soil moisture profile (θ_*S,i,t*_), in turn, then provides a basis to study the impact of other soil textures. A new soil texture specific *Ψ*_*S,i,t*_ profile can then be deduced by using θ_*sat*_, Ψ_*sat*_ and *b* values corresponding to different soil texture types (values from Table 2 of Clapp & Hornberger (1978)). This enabled us to study *Ψ*_*S,i,t*_ profiles for four distinct soil types, i.e. (i) sand, (ii) loam, (iii) sandy clay and (iv) clay soils, in relation with the original silt loam *Ψ*_*S,i,t*_ profile.

**Table 2.** Sampled liana and tree individuals, provided with their species, respective diameter at breast height (DBH, in cm) and their δ^2^H and 910 δ^18^O ranges (in ‰, VSMOW) measured per individual.

| Code | Growth form | DBH [cm] | Family | Species name | $\delta^2\text{H}_x$-range<br>[in ‰, VSMOW] | $\delta^{18}\text{O}_x$-range<br>[in ‰, VSMOW] |
| --- | --- | --- | --- | --- | --- | --- |
| SP1 | Tree | 15.6 | Moraceae | *Coussapoa sp.* | -30.1; -25.5 | -2.8; -2.6 |
| SP2 | Tree | 50.9 | Fabaceae | *Vouacapoua americana* | -23.9; -18.1 | -3.1; -2.2 |
| SP3 | Tree | 44.6 | Vochysiaceae | *Erismia nitidum* | -27.7; -20.8 | -3.2; -1.9 |
| SP4 | Tree | 26.1 | Sapotaceae | *Micropholis guyanensis* | -29.8; -28.0 | -3.0; -2.9 |
| SP5 | Tree | 21.0 | Anacardiaceae | *Tapirira guyanensis* | -31.1; -18.0 | -3.2; -2.2 |
| SP6 | Tree | 49.7 | Fabaceae | *Albizia pedicellaris* | -26.9; -22.1 | -3.2; -2.6 |
| SP1 | Liana | 2.8 | Polygonaceae | *Coccoloba sp* | -27.9; -20.7 | -3.9; -2.3 |
| SP2 | Liana | 2.7 | Convolvulaceae | *sp.* | -29.3; -24.0 | -4.4; -2.9 |
| SP3 | Liana | 0.8 | Moraceae | *sp.* | -40.8; -22.6 | -4.5; -2.3 |
| SP4 | Liana | 3.8 | Combretaceae | *cf. rotundifolium Rich.* | -23.6; -15.2 | -2.9; -2.0 |
| SP5 | Liana | 0.7 | Convolvulaceae | *Maripa cf violacea* | -31.6; -19.7 | -3.8; -2.7 |
| SP6 | Liana | 3.8 | Convolvulaceae | *Maripa sp.* | -35.3; -24.4 | -4.8; -3.1 |

##### Volume of water uptake

We varied the total diurnal volume of water taken up by the tree. New *SF*_*t*_ values are scaled using algorithms from literature that provide an estimate of the daily sap flow volume of a tree based on its DBH (Andrade *et al.*, 2005; Cristiano *et al.*, 2015).

##### Root conductivity

We varied the root membrane permeability (*k*_*R*_) to match multiple species specific values found in literature (Sands *et al.*, 1982; Rüdinger *et al.*, 1994; Steudle & Meshcheryakov, 1996; Leuschner *et al.*, 2004).

The second set of sensitivity analyses test the impact of root hydraulics, sap flow velocities and sampling strategies on the sampled *δ*^*2*^*H*_*X*_. We obtained 1000 samples per parameter from corresponding distributions and ranges (given in Table S2) with a Latin hypercube approach (McKay *et al.*, 1979; McKay, 1988). This is a stratified sampling procedure for Monte Carlo simulation that can efficiently explore multi-dimensional parameter space. In brief, Latin Hypercube sampling partitions the input distributions into a predefined number of intervals (here 1000) with equal probability. Subsequently, a single sample per interval is extracted in an effort to evenly distribute sampling effort across all input values and hence reduce the number of samples needed to accurately represent the parameter space.

### Part B: Empirical exploration

#### Data on variation in i-H_2_O-xyl with plant height

We used data for six canopy trees and six canopy lianas sampled on two subsequent dry days (24-25 August, 2017) at the Laussat Conservation Area in Northwestern French Guiana. The sampling site (05°28.604’N-053°34.250’W) lies approximately 20 km inland at an elevation of 30 m a.s.l. This lowland rainforest site has an average yearly precipitation of 2500 mm yr^−1^ (Baraloto *et al.*, 2011). Average and maximum daily temperature of respectively 30°C and 36°C were measured during the sampling period. Sampled individuals are located in the white sands forest habitat (Baraloto *et al.*, 2011), on a white sandy ultisol with typically high percentage of sand.

Individuals (Table 2) were selected based on assessment of climbable tree, intactness of leafy canopy vegetation and close vicinity with one another to optimize similarity in meteorological and edaphic characteristics. Liana diameters were measured at 1.3 m from the last rooting point (Gerwing *et al.*, 2006), tree diameters were measured at 1.3 m (Table 2). Sampling was performed between 9am and 2pm to assure high sap flow. Liana and tree sampling allowed highly contrasted sap flow velocities (Gartner *et al.*, 1990).

##### Sampling strategy

The stem xylem tissue of individual plants was sampled at different heights (1.3, 5, 10, 15 and 20 m where possible) at the same stem orientation. The order of sampling, i.e. ascending versus descending heights, was randomized. Tree stem xylem samples were collected with an increment borer (5 mm diameter), resulting in wooden cylinders from which bark and phloem tissues were removed. Coring was performed within the horizontal plane at the predefined heights, oblique to the center of the stem to maximize xylem and minimize heartwood sampling, and slowly to avoid heating up the drill head and kinetic fractionation. Taking one sample generally took between 5 and 10 minutes. Since coring lianas was not possible, we collected cross-sections of the lianas after removing the bark and phloem tissue with a knife. All materials were thoroughly cleaned between sampling using a dry cloth to avoid cross-contamination. Upon collection, all samples were placed in pre-weighed glass collection vials, using tweezers, to reduce contamination of the sample. Glass vials were immediately sealed with a cap and placed in a cooling box, to avoid water loss during transportation.

##### Sample processing

Sample processing was performed as in De Deurwaerder *et al.* (2018). Specifically, all fresh samples were weighed, transported in a cooler and frozen before cryogenic vacuum distillation (CVD). Water was extracted from the samples via CVD (4 h at 105°C). Water recovery rates were calculated from the fresh weight, weight after extraction and oven dry weight (48 h at 105°C). Samples were removed from the analysis whenever weight loss resulting from the extraction process was below 98% (after Araguás-Araguás *et al.*, 1998). Sample isotopic signatures were measured by a Wavelength-Scanned-Cavity Ring-Down Spectrometer (WS-CRDS, L2120-i, Picarro, California, USA) coupled with a vaporizing module (A0211 High Precision Vaporizer) through a micro combustion module to avoid organic contamination (Martin-Gomez *et al.*, 2015; Evaristo *et al.*, 2016). Internal laboratory references were used for calibration, with measurement precision of ±0.1‰ and ±0.3‰ for δ^18^O and δ^2^H, respectively. Post-processing was performed using SICalib (version 2.16; Gröning, 2011)

Isotopic composition, expressed in terms of [^18^O]/[^16^O] and [^2^H]/[^1^H] ratios, is represented by *δ*-values (in our case, *δ*^*18*^*O* and *δ*^*2*^*H*), which indicate the deviation from a designated standard (i.e. V-SMOW, Vienna Standard Mean Ocean Water) in parts per thousand (expressed in ‰): 

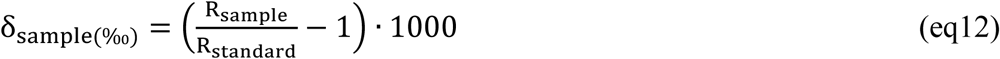

where *R* is the heavy to light isotope ratio measured in the sample or standard. We calculate normalized *i-H*_*2*_*O-xyl* for each individual at every sampled height *h* (*ε*^*2*^*H*_*X*_ and *ε*^*18*^*O*_*X*_) as the bias of each sample from the stem mean (derived from all stem samples of that individual): 

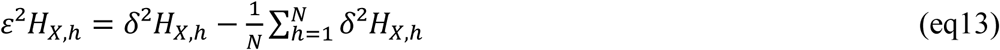

with *N* the number of heights sampled per individual.

#### Data on high temporal-resolution variation in i-H_2_O-xyl

We used data from three extensive field campaigns by Zhao *et al.* (2014) who sampled plant *i-H*_*2*_*O-xyl* at high temporal resolution in the Heihe River Basin (HRB), northwestern China. Four distinct study locations differing in altitude, climatological conditions and ecosystem types were selected. At each location, the dominant tree, shrub and/or herb species was considered for sampling. In August 2009, *Populus euphratica* was sampled in the Qidaoqiao riparian forest (42°01’N-101°14’E) and *Reaumuria soongorica* in the Gobi desert ecosystem (42°16’N-101°17’E; 906-930 m a.s.l). In June–September 2011 *Picea crassifolia, Potentilla fruticose, Polygonum viviparum* and *Stipa capillata* were measured in the Pailugou forest ecosystem (38°33’N-100°18’E 2700-2900 m a.s.l). All species were samples 2-hourly, with the exception of *P. crassifolia* which was measured hourly. Stem samples were collected for trees and shrubs, while root samples were obtained for the herb species (more details in Zhao *et al.* (2014)).

Upon collection, all samples were placed in 8 mL collection bottles and frozen in the field stations before transportation to the laboratory for water extraction via CVD (Zhao *et al.*, 2011). Both *δ*^*18*^*O* and *δ*^*2*^*H* were assessed with an Euro EA3000 element analyzer (Eurovector, Milan, Italy) coupled to an Isoprime isotope ratio mass spectrometer (Isoprime Ltd, UK) at the Heihe Key Laboratory of Ecohydrology and River Basin Science, Cold and Arid Regions Environmental and Engineering Research institute. Internal laboratory references were used for calibration, resulting in measurement precision of ±0.2‰ and ±1.0 ‰ for δ^18^O and δ^2^H, respectively.

## Results

### Part A: Modelling exploration

#### Simulated temporal fluctuation in isotopic signature

##### i. Isotopic signature at stem base and basic model behavior

At the stem base, simulated *δ*^*2*^*H*_*X,0,t*_ displays a diurnal fluctuation (Fig.1) that corresponds to the daily sap flow pattern (Fig.1c). This pattern is caused by shifting diurnal RWU depth. Early in the morning, when transpiration is low, most of the RWU occurs in deeper layers, where soil water potential is less negative and isotopic composition is dominated by depleted deuterium (Fig.S2a,b). As transpiration increases during the day, a significant proportion of RWU is extracted from the drier shallow layers, which have an enriched isotopic composition. In the afternoon, as transpiration declines, isotopic composition reflects again the composition of the depleted deep soil and it remains constant throughout the night because SWIFT does not consider mixing of the internal water in stem and roots nor hydraulic lift.

**Fig 1.**
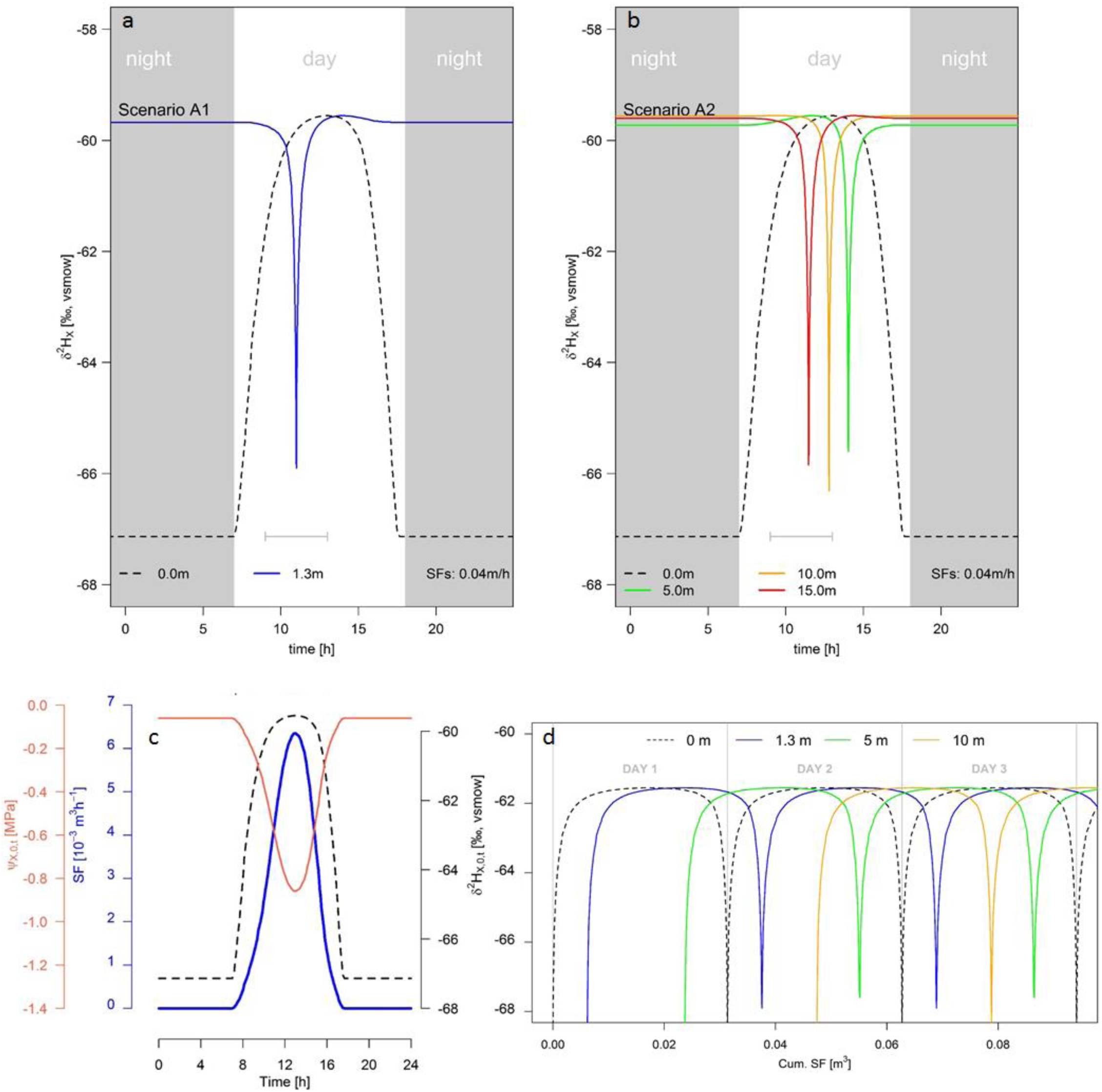
**Panel a & b:** Diurnal patterns of simulated xylem water deuterium signatures (δ^2^H_X_) fluctuation as a function of time for various tree heights. The modeled tree has an average daily sap flow speed (SF_S_) of 0.04 m h^−1^, which corresponds to an average daily true sap flow velocity of 0.28 m h^−1^ (SF_V_), and the standard parameterization is detailed in Table S1. Panel shows analysis A1 output where diurnal δ^2^H_X_ patterns are provided at stem base (0 m, black dashed line) and at general tree coring height at breast level, i.e. at 1.3 m (blue). Panel shows analysis A2 outputs demonstrating diurnal patterns in δ^2^H_X_ within a standard tree at various heights, i.e. at 0 m (black dotted), 5 m (green), 10 m (orange) and 15 m (red). These heights represent random branch sample collection and conform to the standard practice of RWU assessment. Grey lines with whiskers indicate the common sampling period (9:00 until 13:00) according to standard practice. **Panel c:** Sap flow (SF, blue line), xylem water isotopic signature (*δ*^2^*H*_*X*,0,*t*_ black dashed line) and water potential at stem base (*Ψ*_*X*,0,*t*_, red line) are shown over the period of a single day. **Panel d:** Simulated δ^2^H_X_ fluctuations in function of the cumulative sap flow volume measured at various heights: stem base (0 m, black dashed), 1.3 m (blue), 5 m (green) and 10 m (red). Days are delineated by grey vertical lines.

The most enriched *δ*^*2*^*H*_*X*_-values (approx.-59‰) are found in alignment with the diurnal minimum of *Ψ*_*X*,0,*t*_ (approx.-0.85 MPa, Fig.1c). At this moment, Δ*Ψ*_*i,t*_ are maximized, enabling water extracting from the upper and driest soil layers. Most root biomass is located near the surface (cf. Jackson *et al.*, 1995, Fig S2c) and uptake in these layers will result in relatively high contributions to the total RWU.

In contrast, ΔΨ_*i,t*_ are smaller in the early morning and late afternoon causing root water uptake in the upper soil layers to halt. The decreasing ΔΨ_i,t_ translates into higher proportions of RWU originating from deeper, more depleted soil layers. This causes *δ*^*2*^*H*_*X*_ to drop to a baseline of approx. −67‰. This afternoon depletion of *δ*^*2*^*H*_*X*_ will henceforth be indicated as the *δ*^*2*^*H*_*X*_-baseline drop.

##### ii. Isotopic signature at different times, heights and SF_V_

Temporal fluctuation in *δ*^*2*^*H*_*X*_ within a tree at 1.3 m (i.e. the standard sampling height; Analysis A1) and at other potential sampling heights (e.g. branch collection; Analysis A2) are provided in Fig.1a and b, respectively. Both analyses show that fluctuations in *δ*^*2*^*H*_*X*_ depend on the height of measurement and the corresponding time needed to move the water along the xylem conduits. Note that whether the *δ*^*2*^*H*_*X*_-baseline drop at any height reflects the stem base minimum depends on the selected temporal resolution (here 1 min, see Fig.S5). The relation between *δ*^*2*^*H*_*X*_ variance and cumulative sap flow volumes is provided in Fig.1d. Here, the piston flow dynamics in SWIFT originate from lateral translation of the *δ*^*2*^*H*_*X*_ fluctuation at *δ*^*2*^*H*_*X,0,t*_. In addition to sampling height, analysis A3 depicts the importance of sampling time (Fig.2).

**Fig 2.**
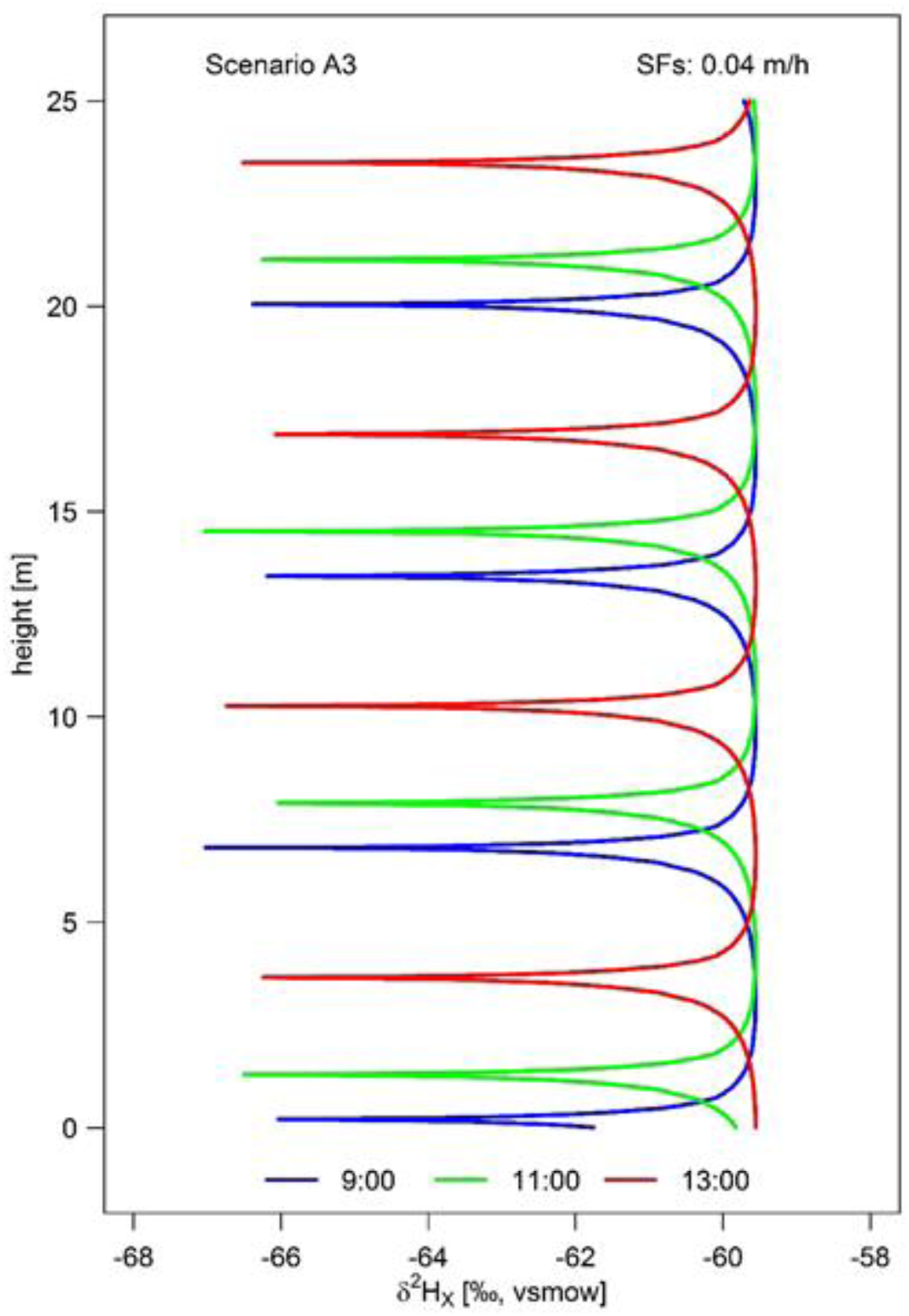
Model outputs for model analysis A3 representing the xylem water deuterium signatures (δ^2^H_X_) as a function of the tree height simulated for different sampling times (9:00 in blue, 11:00 in green and 13:00 in red). The modeled tree has an average daily sap flow speed of 0.04 m h^−1^ (SF_S_), which corresponds to an average daily true sap flow velocity of 0.28 m h^−1^ (SF_V_), and the standard parameterization is detailed in Table S1.

Analysis B outputs predict the occurrence and width of the *δ*^*2*^*H*_*X*_*-*baseline drop as a function of *SF*_*V*_ (Fig.3). Moreover, depending on SF_V_, the isotopic signal can take hours or days to travel from roots to leaves, as observed experimentally (Steppe *et al.*, 2010). Low *SF*_*V*_ allows multiple *δ*^*2*^*H*_*X*_-baseline drops over the length of a single tree. This means that sampled *δ*^*2*^*H*_*X*_ can reflect soil isotopic composition of the past several days. This has direct implications for comparing samples obtained at different times and heights and for species that experienced different *SF*_*V*_ histories.

**Fig 3.**
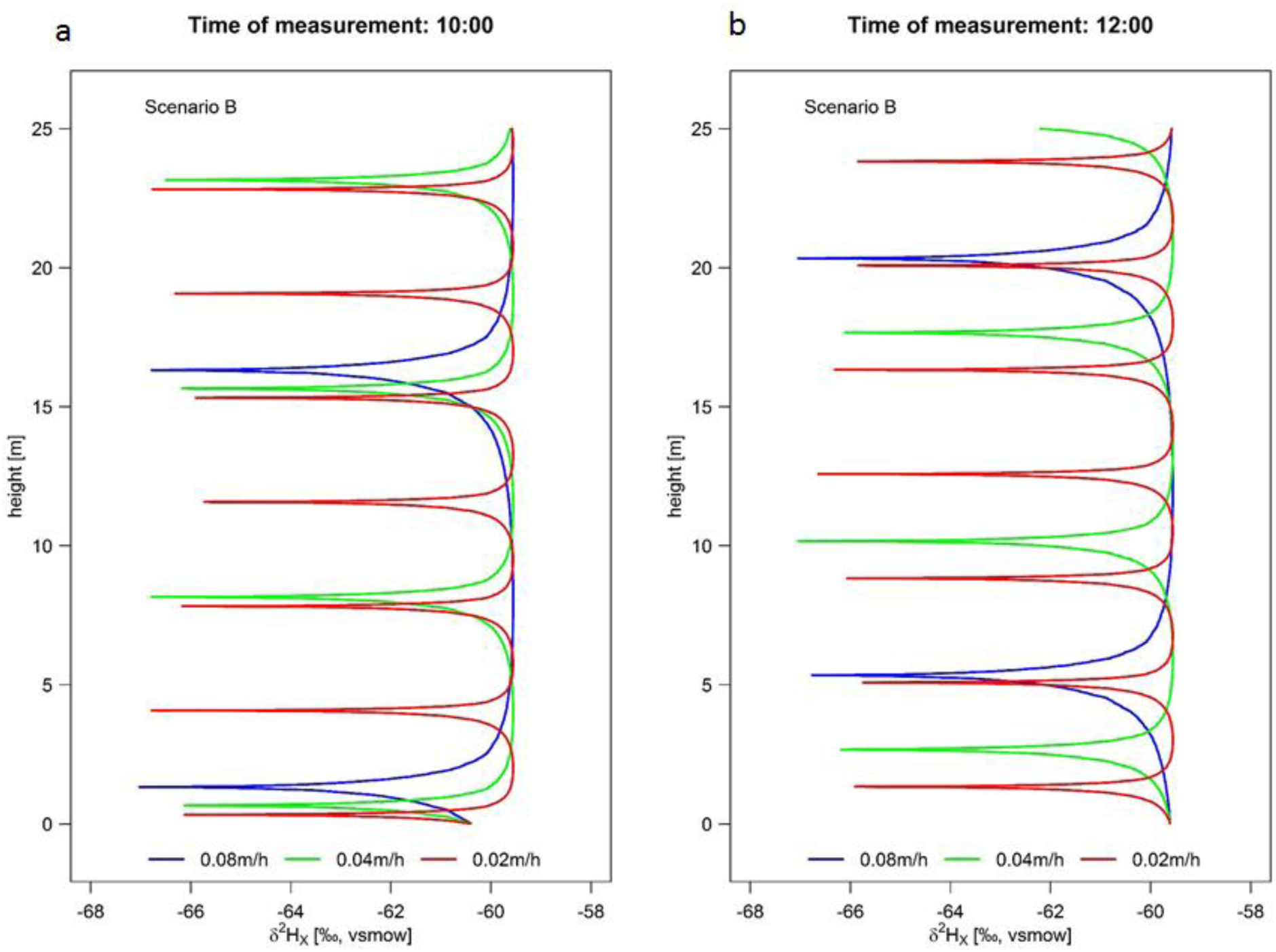
Model outputs for model analysis B representing the xylem water deuterium signatures (δ^2^H_X_) along the tree height at 10:00 h. (a) and 12:00 h. (b). Modeled trees all have the standard parameterization detailed in Table S1, differing only in their average sap flow speed, i.e. 0.08 (blue), 0.04 (green) and 0.02 m h^−1^ (red) (corresponding to an average true sap flow velocity SF_V_ of. 0.56, 0.28 and 0.14 m h^−1^, respectively).

#### Potential biases in root depth estimation

Both timing of measurement (Fig.4a) and *SF*_*V*_ (Fig.4b) influence rooting depth estimates derived via the direct inference and end-member mixing analysis method (Fig.S2) (Rothfuss & Javaux, 2017). Collection of tree samples at 1.30 m can result in erroneous estimation, deviating up to 104% from the average daily RWU depth (Fig.4). Plotting the relative error in RWU depth as a function of time and *SF*_*V*_ (Fig.4c), shows the timing of sampling when *δ*^*2*^*H*_*X*_ measurements at a set height (i.e. at 1.30 m) capture unbiased estimates of the average RWU depth. Xylem water sampling should be timed to capture the *δ*^*2*^*H*_*X*_ that corresponds to water extracted at peak RWU, and the expected sampling time can be derived by considering the time needed for the water to reach the point of measurement. In general, SWIFT predicts that plants with slow *SF*_*V*_ should not be measured during the morning hours, as this results in measuring the preceding days’ absorbed water. In contrast, trees with higher *SF*_*V*_ support earlier sample collection.

**Fig. 4.**
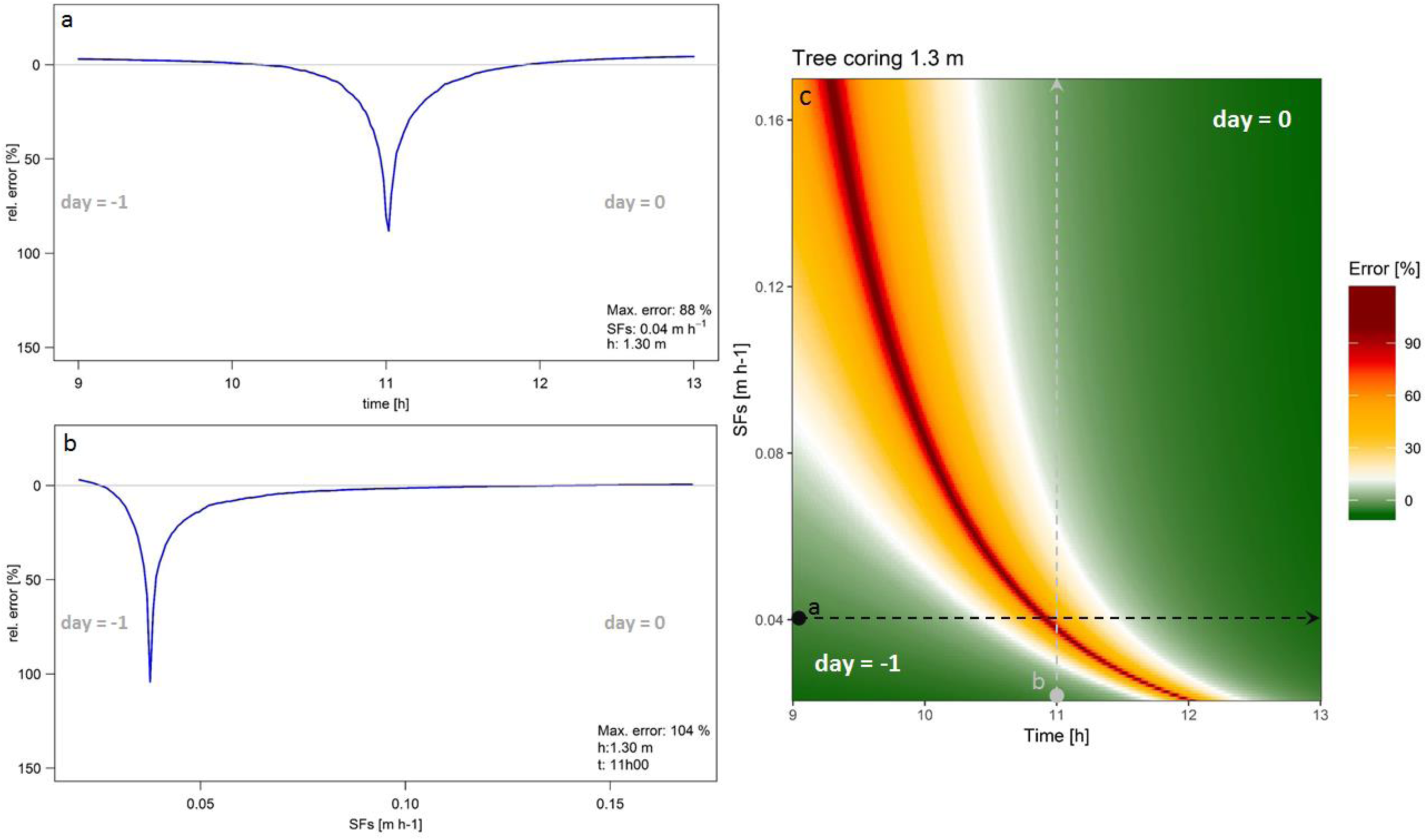
a) Relative error on the inferred root water uptake (RWU) depth (i.e. bias between the average daily and the instantaneous derived RWU depth), for a tree measured at standard tree coring height (i.e. 1.30 m) which has a sap flow velocity (SF_S_) of 0.04 m h^−1^ (i.e. SF_V_ = 0.28 m h^−1^), over the common sampling period (9:00 until 13:00). b) Relative error on the inferred RWU depth considering a tree measured at standard tree coring height (1.30 m) at 11:30, but which differs in SF_S_. c) Relative error on the inferred RWU depth over the duration of the common sampling period (9:00 until 13:00) and over a range of potential SFS (in m h^−1^) – corresponding to SF_V_ range of 0.15–1.25 m h^−1^. Dotted lines a (black) and b (grey) correspond to their respective representation in panel a and b. day= −1 and day= 0 indicate whether the derived RWU depth error corresponds to the previous or current day of measurement.

#### Sensitivity analysis

Our sensitivity analyses shows that the expected absolute error in RWU depth assessment is directly related to both 1) maximum variance in and 2) the probability of sampling non-representative *δ*^*2*^*H*_*X*_ values. The maximum variance depends on the height, while the probability of sampling non-representative areas depends on the width of the “*δ*^*2*^*H*_*X*_-baseline drop” respectively (defined above). Hence, bias in *δ*^*2*^*H*_*X*_ is predominantly a function of the sampling strategy (timing and height of sampling; Fig.S3) in relation to the *SF*_*V*_ of the plant (shown by a strong effect of lumen area and total diurnal RWU volume in Fig.S3) and some biophysical parameter (Fig.5). We summarized the most important variables as predicted by SWIFT, that should be considering in RWU studies below.

**Fig 5.**
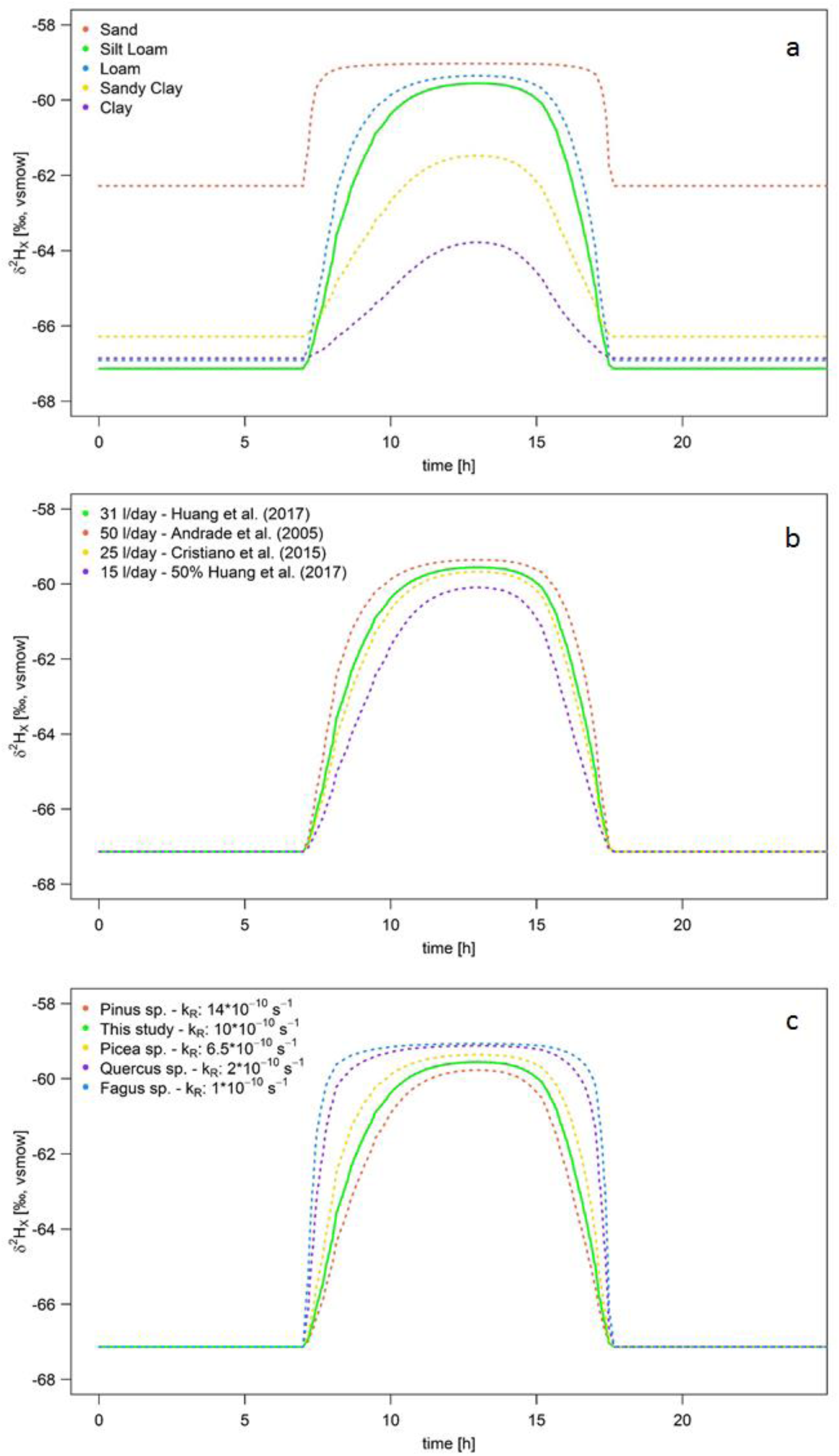
Model sensitivity to (bio)physical parameters. The standard model run is shown by the solid green line in all panels. Panel a: fixed soil moisture and soil deuterium profile (*δ*^2^*H*_*S,i*_) with depth, but with different soil types influencing the soil conductivity and soil water potential gradient in the soil (Ψ_*S,i,t*_). Parameterization for each soil type is derived from Clapp & Hornberger (1978). Panel b: Impact of altering volumes of water taken up by the plant. Panel c: Effect of altering values of root membrane permeability (k_R_) values. Values are species-specific and are derived from literature (Sands *et al.*, 1982; Rüdinger *et al.*, 1994; Steudle & Meshcheryakov, 1996; Leuschner *et al.*, 2004). In each panel all other parameters follow the standard plant parameterization (Table S1).

Plants on loam soils show larger diurnal *δ*^*2*^*H*_*X*_ variances (∼8‰) in comparison with those of clay soils (∼3‰). Larger variances correspond to potentially larger error, but the steeper slope of the *δ*^*2*^*H*_*X*_ curve results in a thinner *δ*^*2*^*H*_*X*_-baseline drop. Hence, loam soil can result in potentially the largest errors but this is mediated by a lower probability of sampling non-representative *δ*^*2*^*H*_*X*_ values during the day.

The volume of water taken up by the plant (*SF*_*t*_; Fig 5b) affects xylem water potential of the plant at stem base 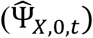. Higher *SF*_*t*_ requires more negative 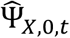 enabling the plant to access more shallow and enriched soil layers. Therefore, an increase in *SF*_*t*_ results in the increase of maximum *δ*^*2*^*H*_*X*_ values (increased maximum error) but also results in a smaller width of the baseline drop (Fig 3). Lower *SF*_*t*_ result in smaller error, but larger probability of sampling an non-representative area (Fig 3).

Root properties, i.e. root membrane permeability (Fig.5c) strongly influence both the total range of *δ*^*2*^*H*_*X*_ variance and the width of the *δ*^*2*^*H*_*X*_-baseline drops. Decreasing root permeability results in thinner *δ*^*2*^*H*_*X*_-baseline drops, but higher maximum *δ*^*2*^*H*_*X*_ variance.

### Part B: Empirical exploration

The observed normalized deuterium signature (*ε*^*2*^*H*_*X*_) along the height of lianas and trees showed strong intra-individual variance exceeding the null model by a factor of 3.2 and respectively (Fig.6a-b). Specifically, differences up to 13.1‰ and 18.3‰ in *δ*^*2*^*H* and 1.3‰ and 2.2‰ in *δ*^*18*^*O* are observed as intra-individual variances for trees and lianas respectively (Table 2).

**Fig 6.**
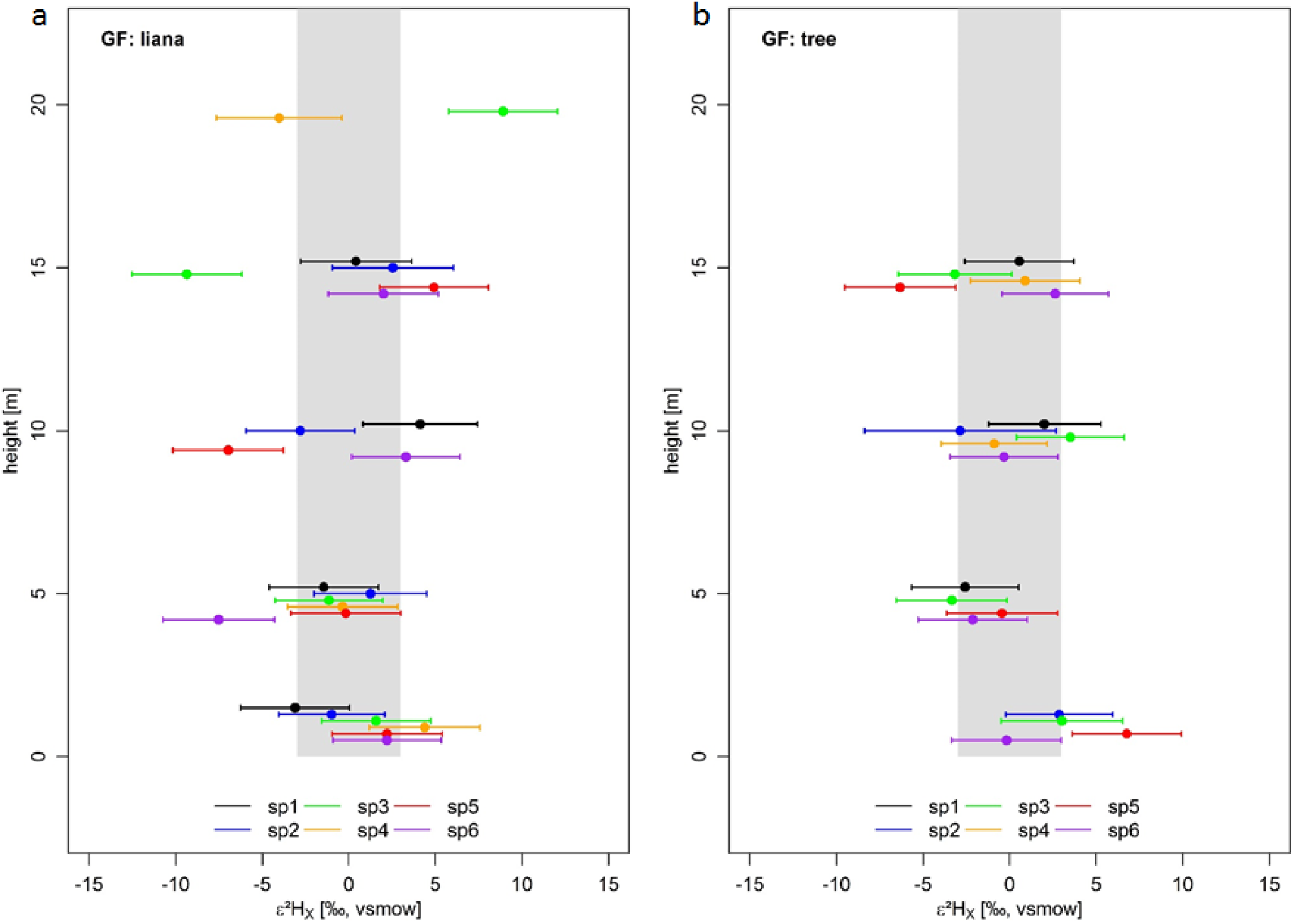
Field measurements of normalized intra-individual δ^2^H_X_ (ε^2^H_X_) for six lianas (panel a) and six trees (panel b). Individuals are provided in different colors; species names can be derived from Table 2. Error whiskers are the combination of potential extraction (± 3‰) and measurement errors of the isotope analyzer. The full grey envelope delineates the acceptable variance from the stem mean (i.e. 3‰) according to the standard assumption of no variance along the length of a lignified plant, i.e the null model.

Similarly, excessive diurnal intra-individual *δ*^*2*^*H*_*X*_ variances emerge in all considered growth forms (Fig.7). Observed maximums were 18.0‰, 21.0‰ and 25.2‰ in *δ*^*2*^*H*_*X*_ for trees, shrubs and herbs respectively (Fig.7; 2.8‰, 6.8‰ and 6.5‰ in *δ*^*18*^*O*_*X*_ in Fig.S4). The null model expected diurnal variance was exceeded for each species during its measurement period, with the exception of *δ*^*2*^*H*_*X*_ measurements of *P. euphratica*. The latter is a riparian forest species, living along the river course, where an easily accessible and abundant ground-water reservoir drives its RWU and *i-H*_*2*_*O-xyl* signal.

**Fig 7.**
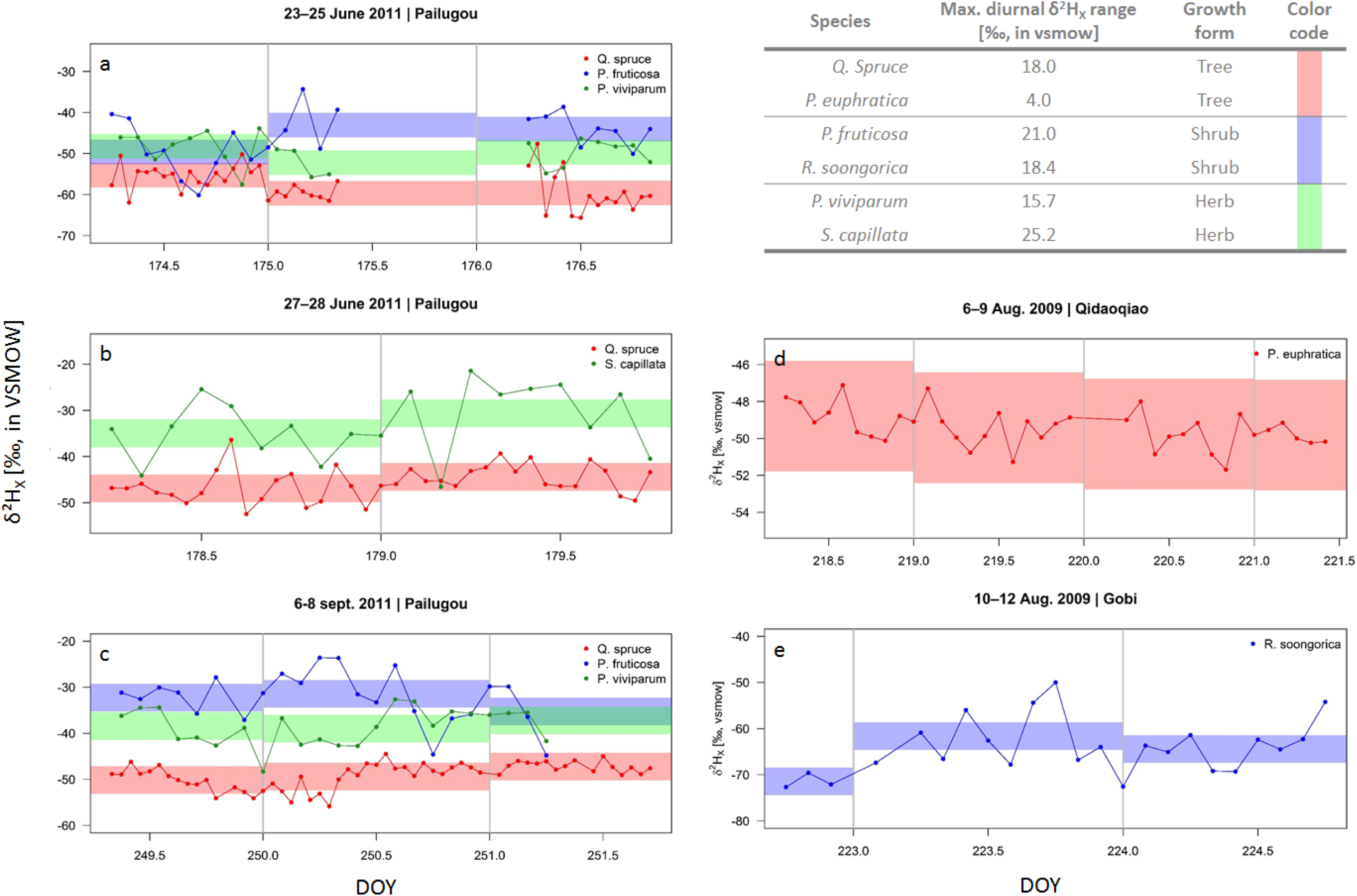
High temporal field measurements of xylem water deuterium isotopic signature (δ^2^H_X_) of two tree (red, stem samples), two shrub (blue, stem samples) and two herb (green, root samples) species sampled in the Heihe River Basin (northwestern China) shown for the respective measurement periods. Timing and location of sampling are provided in the panel titles. The full colored envelope per respective species delineates the acceptable variance from the stem mean (i.e. 3‰) according to the standard assumption of no variance along the length of a lignified plant. Grey vertical lines mark the transition of days. The table provides the maximum measured diurnal δ^2^H_X_ range per species.

## Discussion

### Dynamic diurnal isotopic signatures along plant stems

Our model shows that basic plant hydraulic functioning will result in shifting mixtures of *δ*^*2*^*H*_*X*_ entering the plant (Fig.1a). Daily Ψ_*X*,0,*t*_ fluctuations interact with the Ψ_*S,i,t*_ profile causing different parts of the root distribution to be active during the day. The fluctuations in *δ*^*2*^*H*_*X*_ at the stem base propagate along the xylem with a velocity proportional to the sap flow and this produces variability in sampled *δ*^*2*^*H*_*X*_ that is much larger than the expected measuring error. In addition, empirical field data show excessive *i-H*_*2*_*O-xyl* variance along the stem length (Fig.6) and over a short time frame (i.e. sub-daily, Fig.7). Therefore, the assumption of uniform *δ*^*2*^*H*_*X*_ along the length of a lignified plant is rejected, both theoretically and empirically. Consequently, rather than being static, *δ*^*2*^*H*_*X*_ values along the height of a plant should be considered a dynamic diurnal process.

Importantly, the SWIFT model shows that violation of this assumption results in incorrect assessment of differences in RWU depths between plants. Differences do not necessarily result from variability in RWU depth, but may result from monitoring plants at different heights (Fig.1), at different times (Fig.2) or by comparing individuals which have different *SF*_*V*_ (Fig.3). Our sensitivity analysis reveals that various soil and plant characteristics have an important role in determining both the daily maximum *δ*^*2*^*H*_*X*_ variance as well as the width of the *δ*^*2*^*H*_*X*_-baseline drop. These two characteristics directly impact (i) the expected maximum bias in estimates of RWU depth and (ii) the chance of measuring *δ*^*2*^*H*_*X*_ values that do not represent a mixture of all rooting layers during peak RWU (i.e. the baseline drop). Our work supplements the recent overview of Penna *et al.(* 2018) discussing challenges in using stable water isotopes to study the terrestrial water fluxes. We additionally advocate that future research should explore the minimum set of (bio)physiological drivers and processes that require quantification to correctly interpret *δ*^*2*^*H*_*X*_ along the length of a plant.

### General applicability of model and results

A necessary condition for diurnal shifts in RWU is the existence of a water potential heterogeneity, e.g. more negative water potentials in the upper layers where trees usually have higher root density, which causes a disproportional partitioning of diurnal RWU between deep and shallow roots. Since such a gradient is formed when the upper soil layers undergo evaporation, these conditions are also necessary for the existence of a soil isotopic gradient. Thus, the problem we have identified is intrinsic to the isotopic tracing method for RWU assessment.

Plant transpiration results from complex interaction between atmospheric demands (i.e. driven by VPD and radiation) and stomatal conductance which depends on tolerance of drought stress and soil moisture content. We may expect diurnal fluctuation in radiation and VPD, and hence in water transport and depth of water absorption, as modelled here to be a general phenomenon in nature. Hence, there is a real risk of misinterpretation and calculation errors within the existing literature whenever *i-H*_*2*_*O-xyl* are used to asses RWU and water competition strategies. Moreover, much greater fluctuations in VPD and radiation should be expected under natural conditions than the diurnal cycle described here, and these will increase variability of transpiration fluxes, leading to even more complex dynamics of Ψ_*X*,0,*t*_. For instance, slight alterations in these variables, i.e. a changing degree in cloud cover, can influence Ψ_*X*,0,*t*_ rather abruptly (for e.g. lianas; Chen *et al.*, 2015) and lead to instantaneous changes in *δ*^*2*^*H*_*X*_. Clearly this further complicates the comparison of samples from different plants and sampled at different heights and times, to date overlooked in RWU assessments, and our model certainly illustrates that these considerations are non-trivial.

Note that, based on our model, we expect that soil isotopic enrichment experiments will generate extensive *δ*^*2*^*H*_*X*_ variation along the length of trees whenever diurnal RWU fluctuations cause water extraction to shift between labeled and unlabeled soil layers. Furthermore, when enrichment experiments target trees with different hydraulic properties (such as *SF*_*V*_) care should be taken as to determine when and where to sample these trees in order to assess an enriched signature. Researchers should be certain the signal will be present at the sample height (see e.g. Fig.3).

### Alternative causes of i-H_2_O-xyl fluctuation

The goal of the SWIFT model is to provide a trackable mechanistic explanation using diurnal variations in *SF*_*t*_ and RWU to explain the excessive variance and dynamic nature of the *i-H*_*2*_*O-xyl* fluctuations with plant height and time in field samples (e.g. Fig.7) and elsewhere (Cooper *et al.* 1991). We believe that our model provides a plausible and simple explanation for diurnal *i-H*_*2*_*O-xyl* variation, and that this mechanism contributed to the variation that is observed empirically. Nevertheless, the model necessarily represents a simplified representation of plant hydrological functioning and is therefore limited. There may be alternative causes that contribute to the observed intra-individual *i-H*_*2*_*O-xyl* variances. We discuss these here.

### 1. Fractionation at root level

An increasing body of observations show the occurrence of isotopic fractionation at the root level governed by root membrane transport (Lin & Sternberg, 1993; Vargas *et al.*, 2017) or by unknown reasons (Zhao *et al.*, 2016). Brinkmann *et al.* (2019) hypothesize that root level fractionation causes disparity when RWU depth calculations based on *δ*^*2*^*H*_*X*_ measurements are compared with those of *δ*^*18*^*O*_*X*_. However, it is difficult to imagine a scenario where root fractionation by itself can explain the observed diurnal fluctuations in *i-H*_*2*_*O-xyl* with height and time. Even if root fractionation significantly contributed to variation in *i-H*_*2*_*O-xyl*, we would still need to take into account diurnal fluctuation in RWU to explain the observed patterns.

### 2. Temporal and spatial soil dynamics

The dynamics of soil water movement is complex and soil water content can be extremely heterogeneous in the three spatial dimensions and such variation is currently not represented in SWIFT. Hydraulic lift is a process that generates a vertical redistribution of water in the soil through the roots (Dawson & Ehleringer, 1993), which may change the soil water isotopic composition and mixture drawn up by roots. However, hydraulic lift should redistribute and mix the depleted isotopic signal of deeper layers with the enriched signal of shallower layers. This should lead to lower variation in the soil profile, and less variation along plant length, as such hydraulic lift cannot explain the observed patterns. Heterogeneity in horizontal distribution of water pockets may also affect *i-H*_*2*_*O-xyl* variance. Under these conditions, the horizontal distribution of the absorptive root area becomes more important. However, as the Ψ_*S,i,t*_ and the soil water isotopic signatures of these pockets are interlinked, the mechanistic driver of water extraction is the diurnal fluctuation in water potential gradients in the plant, conform SWIFT.

### 3. Storage tissue and phloem enrichment

Storage tissues release water and sugars in the xylem conduits on a daily basis to support water transpiration demand (Goldstein et al., 1998; Morris et al., 2016; Secchi et al., 2017) or to repair embolism (Salleo et al., 2009; Secchi et al., 2017). Both water and sugars are transported in and out storage tissue via symplastic pathways using plasmodesmata and aquaporins (Knipfer et al., 2016; Secchi et al., 2017), a pathway which has been linked to isotopic fractionation in roots (Ellsworth & Williams, 2007). Moreover, phloem transports photosynthetic assimilates constructed at the leaf level potentially affected by transpiration fractionation (Gessler *et al.*, 2013). Hence, these metabolic molecules might be constructed from enriched ^*2*^*H* and ^*18*^*O* atoms. Water release from storage or phloem tissue might locally alter *i-H*_*2*_*O-xyl* (White et al., 1985). Additionally, time between water storage and release could bridge multiple days, and corresponding isotopic signatures may reflect different soil conditions. It is evident that such dynamics are complex, and it is hard to predict how storage tissue and phloem enrichment affect the *i-H*_*2*_*O-xyl* patterns observed here. Xylem isotopic sampling cannot differentiate between water resulting from RWU or storage, and therefore we cannot discount the possibility that tissue and phloem enrichment play a role. At a minimum this adds further uncertainty to RWU assessment.

Further studies should determine whether the implementation of additional mechanisms such as tree capacitance, root and stem level fractionation, spatiotemporal soil water dynamics, more detailed root systems or storage tissues impact the intra-individual *i-H*_*2*_*O-xyl* and should be accounted for to improve RWU assessment and interpretation

### The way forward

Combining a plant hydraulic model with *in situ SF*_*V*_ and *in situ* Ψ_*S,i,t*_ can help improve the robustness of RWU assessment and interpretation. Measurements of Ψ_*S,i,t*_ at multiple depths should be especially valuable since the SWIFT model showed high sensitivity to alterations of this variable and these can be directly supplied as model inputs. At the same time, the availability of *SF*_*t*_ measurements allows for identifying the moment when water uptake from all root layers is at its maximum, which can be used to determine the optimal timing of sampling at a given height providing a more robust estimation of RWU depth and uptake.

Alongside the modeling and theoretical approach presented here, new ways to study *δ*^*2*^*H*_*X*_ at a high temporal scale are strongly encouraged. For example, pioneering work of Volkmann *et al.* (2016) to the development of an *in situ* continuous isotope measurement technique that offers the possibility for monitoring *i-H*_*2*_*O-xyl* at a sub hourly resolution. This technique holds strong promises for further elucidating the natural *δ*^*2*^*H*_*X*_ variances found within plants and the physiology processes from which these variances result. Such high temporal resolution of isotope measurements, coupled with *in situ* monitoring of various environmental and plant biophysical metrics, are needed for both model improvement and further validation. Moreover, these seem inevitable to eventually differentiate all causal mechanisms of the observed intra-individual *i-H*_*2*_*O-xyl* variance.

## Conclusions

We have demonstrated that the assumption of no intra-individual *i-H*_*2*_*O-xyl* variation is rapidly violated once models incorporate even basic plant hydraulic functioning. Moreover, the incorrectness of this assumption is confirmed by empirical field data, showing excessive variance and high temporal fluctuations in *i-H*_*2*_*O-xyl*. We expect the observed *i-H*_*2*_*O-xyl* variance and sub-daily fluctuations result, in part, from the mechanisms considered in the SWIFT model, though they likely represent an end product of various physiological processes which impact *i-H*_*2*_*O-xyl*.

Our theoretical explorations warn that variability in the isotopic signature can result in erroneous RWU depth estimation and will complicate the interpretation and comparison of data: samples taken at different heights, times or plants differing in *SF*_*V*_ may incorrectly show differences in RWU depth. We further predict that various soil parameters and plant hydraulic parameters affect (i) the absolute size of the error and (ii) the probability of measuring *i-H*_*2*_*O-xyl* values that do not represent the well-mixed values during the plants’ peak RWU. Hydraulic models, such as SWIFT, should be used to design more robust sampling regimes that enable improved comparisons between studied plants. We advocate to add *SF*_*t*_, which indirectly reflects diurnal RWU fluctuations, and Ψ_S,i,t_ monitoring as a minimum in future RWU assessments since these parameters were predicted to be the predominant factors introducing variance in *i-H*_*2*_*O-xyl* from the SWIFT model exploration. However, soil texture and root permeability are also key considerations to measure especially when comparing across species and sites.

Our findings do not exclude additional factors that impact the observed intra-individual *i-H*_*2*_*O-xyl* variance and temporal fluctuation. Therefore, we strongly emphasize the need for more testing. Directed studies that validate and quantify the relative impact of other plant physiological processes towards variance in *i-H*_*2*_*O-xyl* are a prerequisite before improved modeling tools can be developed, and bias in RWU assessments eliminated.

## Acknowledgement

This research was funded by the European Research Council Starting Grant 637643 (TREECLIMBERS), the FWO grants (1507818N, V401018N to HDD), the Carbon Mitigation Initiative at Princeton University (MD, MDV), Agence Nationale de la Recherche “Investissement d’Avenir” grant (CEBA: ANR-10-LABX-25-01), the Belgian American Educational Foundation (BAEF to FM) and the WBI (FM). We are grateful to Samuel Bodé, Megan Bartlett, Isabel Martinez Cano and Pedro Hervé-Fernández who provided feedback on analytical and interpretative aspects of the study. We thank Dries Van Der Heyden, Wim Van Nunen, Laurence Stalmans, Oscar Vercleyen, Katja Van Nieuland, Stijn Vandevoorde and Clément Stahl for data collection and lab processing. We credit Pascal Petronelli and Bruce Hoffman for species identification, and Cora N. Betsinger for proofreading. Cheng-Wei Huang’s work provided inspiration for this research.

## Author contribution

H.V., M.D.V and P.B. supervised and provided guidance throughout all aspects of the research. H.D.D., M.D.V and H.V. designed the study. H.D.D., L.Z. and L.W. collected the samples and data during the field campaign and performed the processing and analysis of the samples. The model was developed and coded by H.D.D, M.D.V, M.D. and F.M. All authors contributed to interpretation of the results and to the text of the manuscript.

---

**Fig S1.**
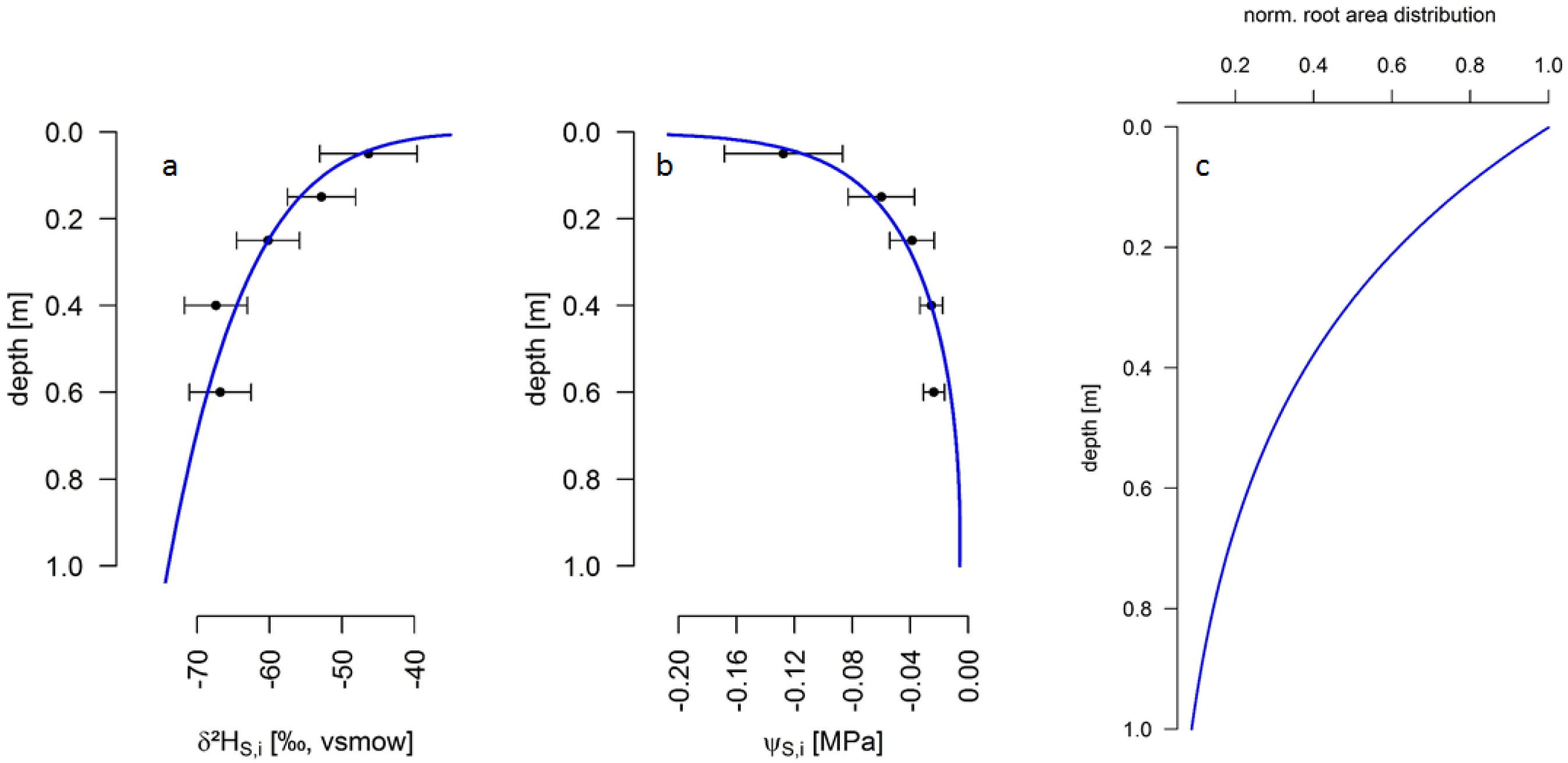
Panel a: Soil water deuterium (δ^2^H_S,i_) signature profile over the soil depth, data from Meißner et al. (2012). Panel b: Soil water potential (Ψ_S,i_) over the soil depth, data from Meißner et al. (2012). Panel c: The fine root surface area distribution with soil depths adapted from Jackson et al. (1995) and normalized to the topsoil. All equations and corresponding parameters for the fitted curves can be found in Table S1.

**Fig S2.**
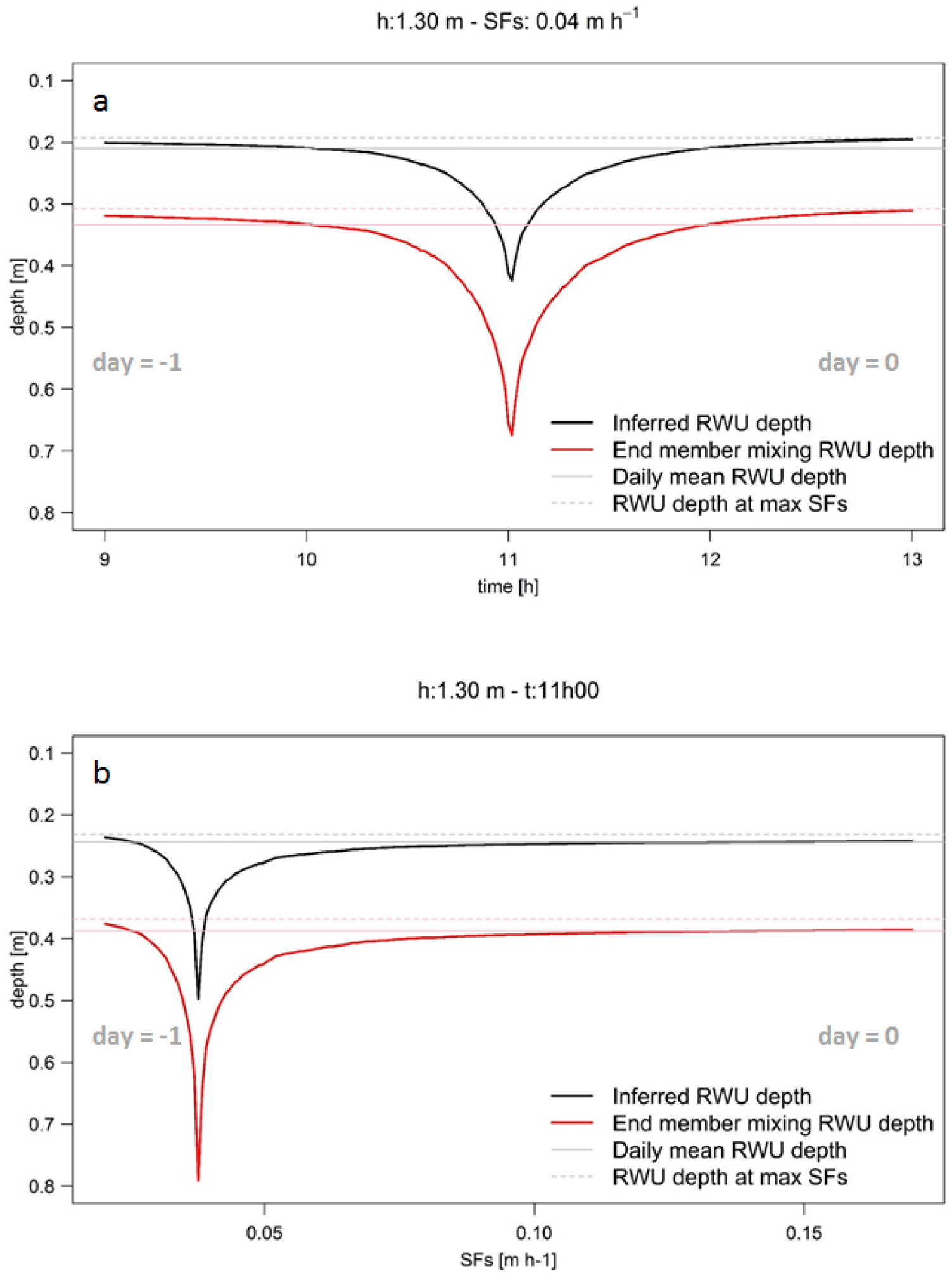
Differences between the root water uptake (RWU) depth derived from using either the direct inference (black line) or the end member mixing (red line) approach. Panel a: The derived RWU depth for a tree sampled at standard tree coring height (i.e. 1.30 m) having a sap flow velocity (SF_S_) of 0.04 m h^−1^ (i.e. SF_V_ = 0.28 m h^−1^), over the common sampling period (9:00 until 13:00). Panel b: The derived RWU depth considering a tree sampled at standard tree coring height (1.30 m) at 11:30, but which differs in SF_S_. The grey and pink solid lines represent daily mean RWU depth while the grey and pink dashed lines represent the RWU depth at peak sap flow activity, respectively, for the direct inference and end member mixing model approach. day= -1 and day= 0 indicate whether the derived RWU depth error corresponds to the previous or current day of measurement.

**Fig S3.**
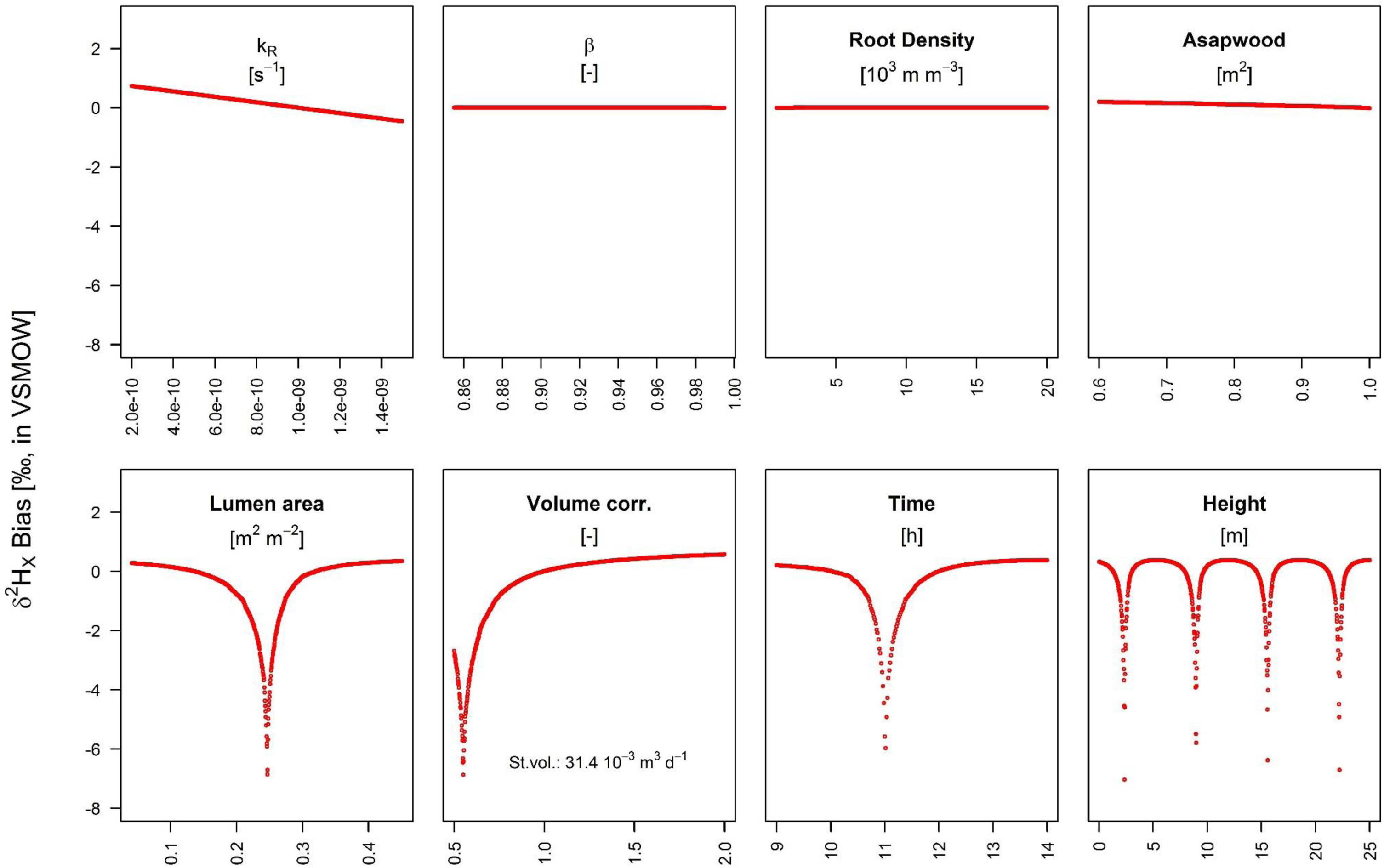
Sensitivity analysis where all parameters are varied one-at-the-time as compared to the standard parameterization (see Table S1). For each studied variable, 1000 model runs were performed, studying the resulting δ^2^H_X_ bias in comparison with the standard run. Each time, the studied parameter value was assigned randomly from a defined probability distribution or range using a Latin Hypercube scheme (see Table S2). Together the root membrane permeability (k_R_, in s^−1^), the β (-) and root density (in 10^3^ m m^3^) form an informative proxy for the soil to root resistance. The lumen fraction (in m^2^ m^−2^), sapwood area (Asapwood, in m^2^) and the total diurnal transported sap flow volume, i.e. net root water uptake (Volume corr., factor of standard run volume), provide an informative proxy for the sap flow velocity. (see Table S1). Time (in h) and height (in m) respectively represent the timing of sampling and the height of sample collection.

**Fig S4.**
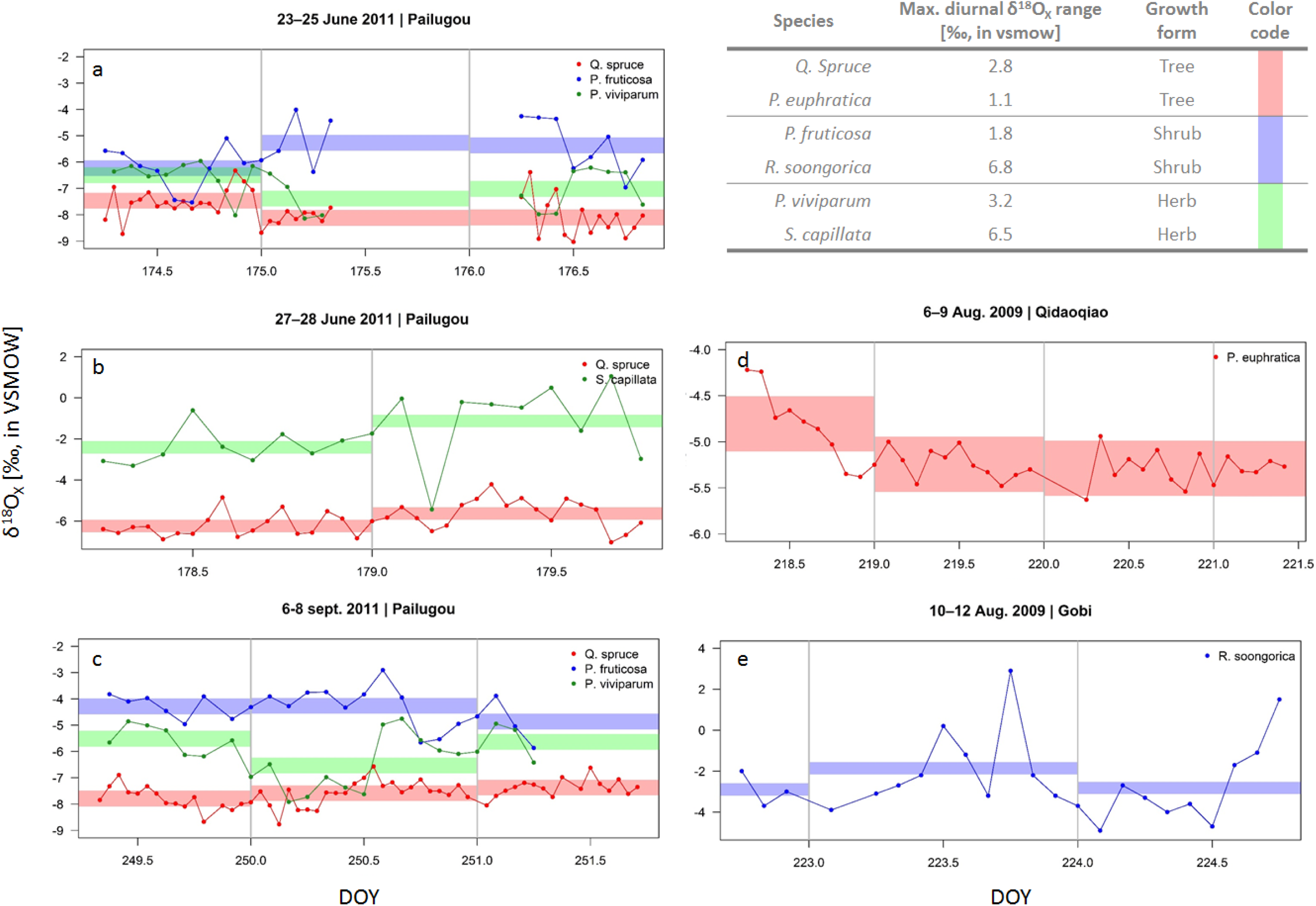
High temporal field measurements of xylem oxygen water isotopic signature (δ^18^O_X_) of two tree (red, stem samples), two shrub (blue, stem samples) and two herb (green, root samples) in the Heihe River Basin (northwestern China) shown for the respective measurement period. Timing and location of sampling are provided in the panel title. The full colored envelope per respective species delineates the acceptable variance from the stem mean (i.e. 0.3‰) according to the standard assumption of no variance along the length of a lignified plant. Grey vertical lines mark the transition of days. The table provides the maximum measured diurnal δ^18^O_X_ range per species.

**Fig S5.**
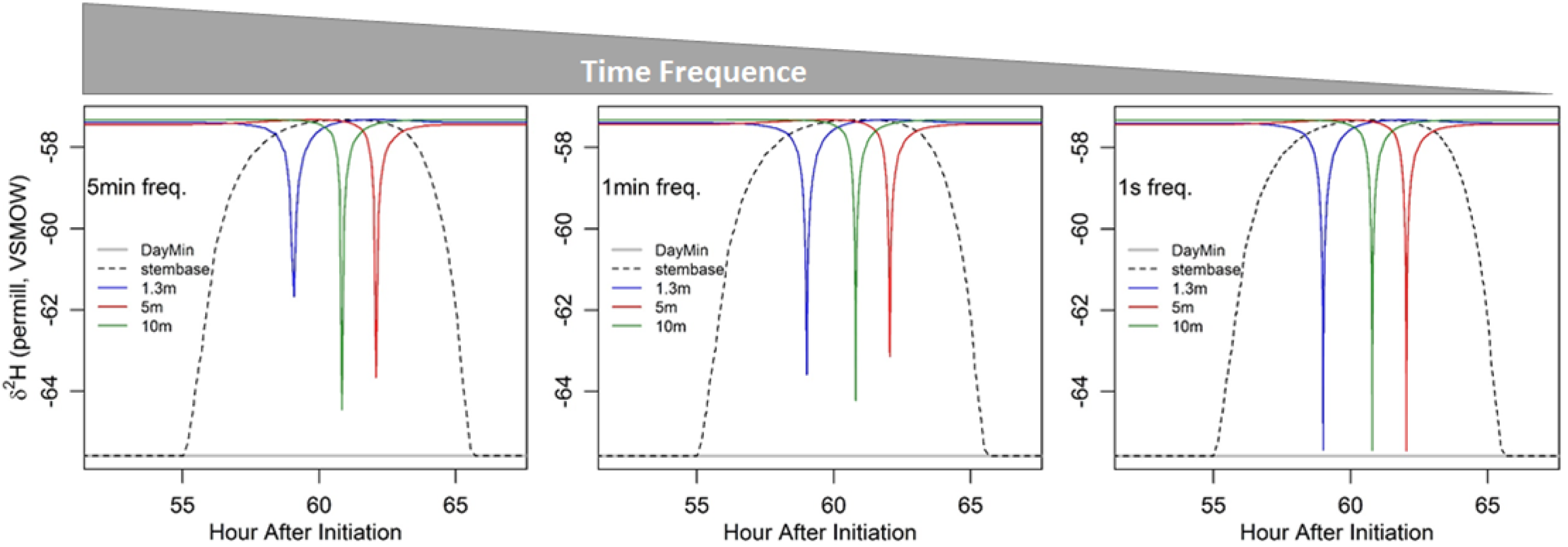
Model simulations performed with varying temporal resolutions, i.e. 5min, 1min and 1sec.

**Table S1.** An overview of the model standard parameterization of the present model, including sap flow, with corresponding references to literature.

| Abbr. | Parameter | Unit | Value | Source |
| --- | --- | --- | --- | --- |
| $A_{Rtot}$ | The plants' total active fine root surface area | $m^2$ | $e^{0.88 \cdot \ln\left(\pi \cdot \left[\frac{DBH \cdot 10^2}{2}\right]^2\right) - 2}$ | Čermák *et al.* (2006)<br>$A_{Rtot} = 23.825 m^2$ |
| $A_{R,i}$ | is the fine root surface area distributed over soil layer $i$. | $m^2$ | $A_{Rtot} \cdot \beta^{100 \cdot z_i} \cdot (1 - \beta^{100 \cdot \Delta z})$ | $A_{Rtot}$ multiplied by the integrated root distribution of each soil layer adapted from Jackson *et al.* (1996) |
| | | | $\beta = 0.976$ | Huang *et al.* (2017) |
| $A_{SAPWOOD}$ | Sapwood area | $m^2$ | $\frac{1.582 \cdot [DBH \cdot 10^2]^{1.764}}{10^4}$ | Meinzer *et al.* (2001) |
| $A_x$ | Total lumen area | $m^2$ | $LF \cdot A_{SAPWOOD}$ | |
| $B_i$ | The overall root length density per unit of soil, not necessarily limited to the studied plant. | $m m^{-3}$ | $R_0 \cdot \beta^{100 \cdot z_i} \cdot \ln(\beta)$ | Adapted from Huang *et al.* (2017)<br>$R_0 = -438\ 688$<br>$\beta = 0.976$ |
| DBH | Diameter at breast height | m | 0.213 | Huang *et al.* (2017) |
| $\delta^2H_{S,i}$ | Soil water isotopic signature of the sampled soil layers | in ‰,<br>VSMO<br>W | $a + (z_i + b)^c$ | Adapted from Meißner *et al.* (2012)<br>...a: -73.98008<br>...b=0.001<br>...c=0.148735; |
| $\Delta z$ | The thickness of each soil layer | m | 0.001 | |
| $f_t$ | Temporal resolution | s<sup>-1</sup> | 1/60 | |
| $k_R$ | Root membrane permeability | s<sup>-1</sup> | 10<sup>-9</sup> | Huang *et al.* (2017) |
| $K_{S,i}$ | The soil hydraulic conductivity defined per soil depth | m s<sup>-1</sup> | $K_{S,max} \cdot \left( \frac{\Psi_{sat}}{\Psi_{S,i,t}} \right)^{2+\frac{3}{b}}$<br>$K_{S,max} = 7.2 \cdot 10^{-6} \text{ m s}^{-1}$<br>$\Psi_{sat} = -0.786 \text{ m H}_2\text{O}$<br>$b = 5.30$ | Huang *et al.* (2017)<br>Clapp & Hornberger (1978)<br>[Table 2, silt loam soil]<br>Clapp & Hornberger (1978)<br>[Table 2, silt loam soil]<br>Clapp & Hornberger (1978) [Table 2,<br>silt loam soil] |
| $LF$ | Lumen fraction per unit sapwood area | $\text{m}^2 \text{m}^{-2}$ | 0.136 | Zanne *et al.* (2010) [Table 2] |
| $SF_t$ | Instantaneous sap flow at time $t$ | $\text{m}^3 \text{s}^{-1}$ | | Adapted from Huang *et al.* (2017)<br>[derived from scenario 6, day 11] |
| $\Psi_{S,i,t}$ | Water potential at a specific soil layer depth $i$ and time $t$ | $\text{m H}_2\text{O}$ | $(a + b \cdot \log(z_i) - c \cdot z_i^2) \cdot CT$ | Adapted from Meißner *et al.* (2012)<br>a: $19.8455 \cdot 10^{-3}$<br>b: $44.8909 \cdot 10^{-3}$<br>c: $25.5594 \cdot 10^{-3}$<br>CT: 101.97 (i.e. conversion factor between MPa and m H<sub>2</sub>O) |

$z_i$ the soil depth of the $i^{\text{th}}$ soil layer (in m)

**Table S2.**
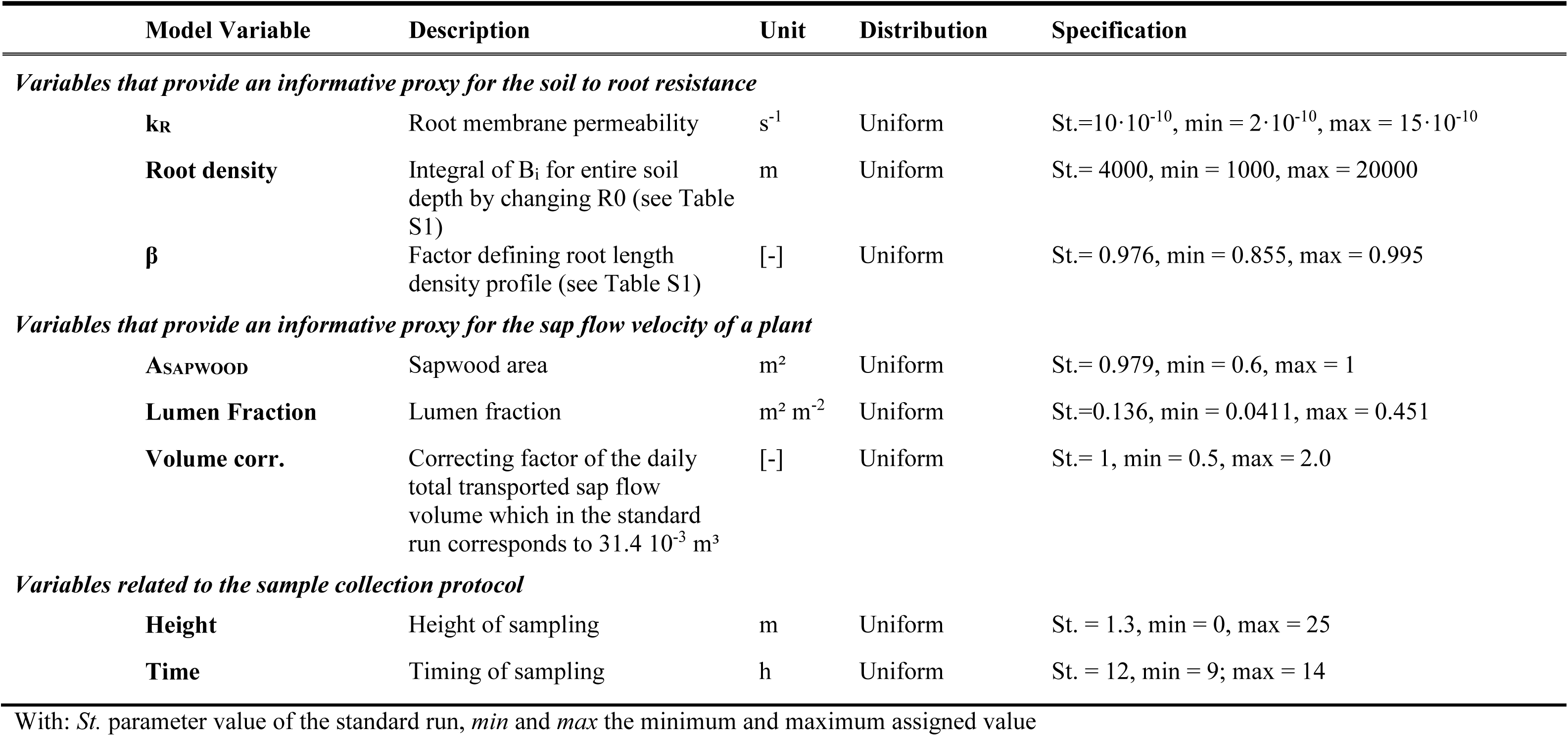
An overview of the defined distribution and ranges used for the sensitivity analysis whose results are displayed in Fig S4.

| Model Variable | Description | Unit | Distribution | Specification |
| --- | --- | --- | --- | --- |
| *Variables that provide an informative proxy for the soil to root resistance* |  |  |  |  |
| **k<sub>R</sub>** | Root membrane permeability | s<sup>-1</sup> | Uniform | St.=10·10<sup>-10</sup>, min = 2·10<sup>-10</sup>, max = 15·10<sup>-10</sup> |
| **Root density** | Integral of B<sub>i</sub> for entire soil depth by changing R0 (see Table S1) | m | Uniform | St.= 4000, min = 1000, max = 20000 |
| **β** | Factor defining root length density profile (see Table S1) | [-] | Uniform | St.= 0.976, min = 0.855, max = 0.995 |
| *Variables that provide an informative proxy for the sap flow velocity of a plant* |  |  |  |  |
| **A<sub>SAPWOOD</sub>** | Sapwood area | m<sup>2</sup> | Uniform | St.= 0.979, min = 0.6, max = 1 |
| **Lumen Fraction** | Lumen fraction | m<sup>2</sup> m<sup>-2</sup> | Uniform | St.=0.136, min = 0.0411, max = 0.451 |
| **Volume corr.** | Correcting factor of the daily total transported sap flow volume which in the standard run corresponds to 31.4 10<sup>-3</sup> m<sup>3</sup> | [-] | Uniform | St.= 1, min = 0.5, max = 2.0 |
| *Variables related to the sample collection protocol* |  |  |  |  |
| **Height** | Height of sampling | m | Uniform | St. = 1.3, min = 0, max = 25 |
| **Time** | Timing of sampling | h | Uniform | St. = 12, min = 9; max = 14 |

With: *St.* parameter value of the standard run, *min* and *max* the minimum and maximum assigned value

